# Decoding the Mechanism of Action of a Parasite TGFβ antagonist Inspires the Creation of Cell-type-specific TGFβ Modulators

**DOI:** 10.64898/2026.02.16.706112

**Authors:** Maarten van Dinther, Tristin Schwartze, Jiying Zhang, Kun Fan, Gerard van der Zon, Luke Power, Cynthia Hinck, Claire Ciancia, Ananya Mukundan, Roman Gonzalez-Prieto, Peter van Veelen, Rick M. Maizels, Andrew P. Hinck, Peter ten Dijke

## Abstract

*Heligmosomoides polygyrus*, a mouse parasite, modulates host immunity by secreting modular transforming growth factor-β (TGFβ) mimics (TGMs). The agonist TGM1 interacts with TGFBR1, TGFBR2, and the co-receptor CD44 through domains D1/2, D3, and D4/5, respectively. In contrast, the antagonist TGM6, which lacks D1/2, but retains TGFBR2 binding through D3, targets different subsets of cells compared to TGM1. The TGM6 co-receptor is unknown. Using X-ray crystallography and binding studies, we show that TGM6 preferentially binds mouse TGFBR2 over human TGFBR2, and that this is essential for its antagonistic function. We identified low-density lipoprotein receptor-related protein 1 (LRP1) and betaglycan (TGFBR3) as co-receptors for TGM6. LRP1 enhances TGM6 efficacy and is vital for its specific antagonistic effects by promoting TGFBR2 degradation, while betaglycan counteracts TGM6 in a TGFBR2-dependent manner. The modular organization of TGMs enabled us to rationally design TGM1/6 chimeras or TGM-D3 fusion with an affibody that recognizes a specific cell-surface receptor, thereby altering cell-type specificity and functionality. Furthermore, we developed a TGFBR2 nanobody that, on its own, has no inhibitory effect but, when fused to a receptor antibody, antagonizes TGFβ signaling in a cell-selective manner. Thus, we designed programmable agents that modulate TGFβ signaling only in target co-receptor-expressing cells.

## 1. Introduction

Transforming growth factor-β (TGFβ) is a secreted cytokine that exerts many diverse cellular responses via specific cell surface heteromeric complexes of two ubiquitously expressed TGFβ receptors, i.e., TGFBR1 and TGFBR2 [1–3]. TGFBR1 and TGFBR2 are essential for TGFβ signaling *(4).* TGFBR1 acts downstream of TGFBR2 [4], and when activated, TGFBR1 induces the phosphorylation of receptor-regulated (R) SMAD2 and SMAD3 proteins [1–3]. Upon activation, R-SMADs form heteromeric complexes with SMAD4, which accumulate in the nucleus and regulate specific gene transcriptional responses [5].

TGF-β plays a pivotal role in physiological processes, including embryonic development, wound healing, and tissue remodeling, as well as in maintaining immune tolerance and preventing autoimmunity. Dysregulated, overactive TGFβ signaling can contribute to pathological responses, such as fibrosis and cancer [1]. However, it is often unclear which critical cell types the multifunctional TGF-β acts on in complex physiological and pathological processes. The same holds for results from studies using current TGFβ inhibitors, such as TGFβ-neutralizing antibodies, TGFBR2-extracellular domain (ECD)-derived ligand traps, and small-molecule TGFBR1 kinase inhibitors, which inhibit TGFβ responses in all cell types [6]. For example, the underlying mechanism by which the TGFβ antagonist inhibits cancer progression is often unclear: does it antagonize cancer cell invasion, inhibit activation of cancer-associated fibroblasts, or inhibit immune evasion and stimulate immune cell infiltration into tumors?

The murine helminth parasite *Heligmosomoides polygyrus* (Hp) modulates host immune responses by secreting TGFβ mimics (TGMs) [7]. The prototypic TGM1 exerts TGF-β-like effects on immune cells in vitro [8–10] and in vivo [8,10–15]. In addition to TGM1, 9 different structurally related HpTGMs have been identified [16]. HpTGMs have no primary sequence similarity to mammalian TGFβ. In contrast to the compact globular protein structure of mammalian TGFβ, TGMs have a modular structure with individual domains that are distantly related to the Sushi domain family or to complement control proteins (CCPs) [17]. Its prototypic member, TGM1, phenocopies the immunosuppressive effects of TGFβ by stimulating T regulatory cells [8]. TGM1 has five separate domains (D) in which D1/2, D3, and D4/5 interact with TGFBR1, TGFBR2, and co-receptor CD44, respectively [18]. CD44-deficient T regulatory cells exhibited impaired TGM1-mediated induction of the transcription factor FoxP3 (*18*). TGM6 lacks D1/2 and, therefore, TGFBR1 engagement, but retains TGFBR2 interaction via its D3. TGM6 is a potent antagonist of TGFBR signaling in fibroblasts, but not splenic T cells [19]. D4/5 of TGM6 is essential for antagonism of TGFβ, but does not bind CD44 [19]. D4/5 is the most divergent among the TGM domains [16], and emerging studies indicate that it is a key determinant of how different TGMs elicit cell-specific effects [18–20].

TGM6 is secreted by a parasite that infects mice, not humans, and has evolved through convergent evolution [21]. Here, we report the molecular mechanisms underlying the selective action of TGM6 on mouse but not human cells. Importantly, we identified low-density lipoprotein receptor-related 1 (LRP1) and betaglycan (TGFBR3) as two opposing TGM6 co-receptors. Moreover, the modular organization of TGMs enabled us to rationally design TGM1/6 chimeras or a fusion of TGM6-D3 with an affibody that recognizes a specific cell-surface receptor, thereby altering the cell-type specificity of TGM6 and transforming the TGM6 antagonist into a mouse-cell-type-selective agonist or antagonist. Furthermore, we developed a TGFBR2 nanobody that, on its own, has no inhibitory effect but, when fused to a receptor antibody, antagonizes TGFβ signaling in a human cell-selective manner. Our identified principles may be instrumental in developing therapies targeting the highly pleiotropic TGFβ in patients with dysregulated TGFβ signaling.

## 2. Results

### 2.1. TGM6-mediated inhibition of TGFβ signaling is effective on mouse and rat, but not human, cells

TGM6 mimics the interaction of TGFβ with TGFBR2, and antagonizes TGFβ signaling in specific cells, including fibroblasts, but not other cell types, such as T cells [19], in a manner dependent on two C-terminal co-receptor-binding domains, D4/5. Although TGM6 D4/5 differs from TGM1 D/5 in that it does not bind CD44 [19], the identity of TGM6 co-receptors remains unknown. In our survey of TGM6-responsive cell lines to select appropriate cells for TGM6 co-receptor identification and mechanism-based studies, we found that TGM6 potently blocked the TGFβ/SMAD signaling response in mouse and rat cells, but not in human cells (Figure 1A). In none of the cells did we observe TGFβ agonistic activity by TGM6 alone (Figure 1A). TGM6 was highly effective in blocking TGFβ-induced SMAD3/4 transcriptional response in mouse NIH3T3 fibroblasts (Figure 1A), and these cells were chosen for further investigation. Analyzing the kinetics of TGM6 inhibition on TGFβ in NIH3T3 cells showed that TGM6 could mitigate TGFβ/SMAD2 phosphorylation after 1 h to 8 h stimulation time (Figure 1B). Consistent with the lack of an effect of TGM6 on TGFβ-induced transcriptional response in human 293T cells, TGM6 did not affect TGFβ-induced SMAD2 phosphorylation (Figure 1C). Increasing the TGM6 doses effectively inhibited TGFβ-induced SMAD3/4 transcriptional response in NIH3T3 cells, in a sharp threshold effect with 10 ng/ml TGM6 being barely suppressive and 25 ng/ml highly suppressive (Figure 1D). Adding TGM6 before the addition of TGFβ was more efficient than adding it at the same time or shortly thereafter (Figure 1E). TGFβ is known for being a potent inducer of epithelial-to-mesenchymal transition (EMT) [22]. TGM6 blocked TGFβ-induced EMT of mouse NMuMG breast cells. Taken together, our results indicate that TGM6 is a potent antagonist of TGFβ in rodent but not human cells.

**Figure 1.**
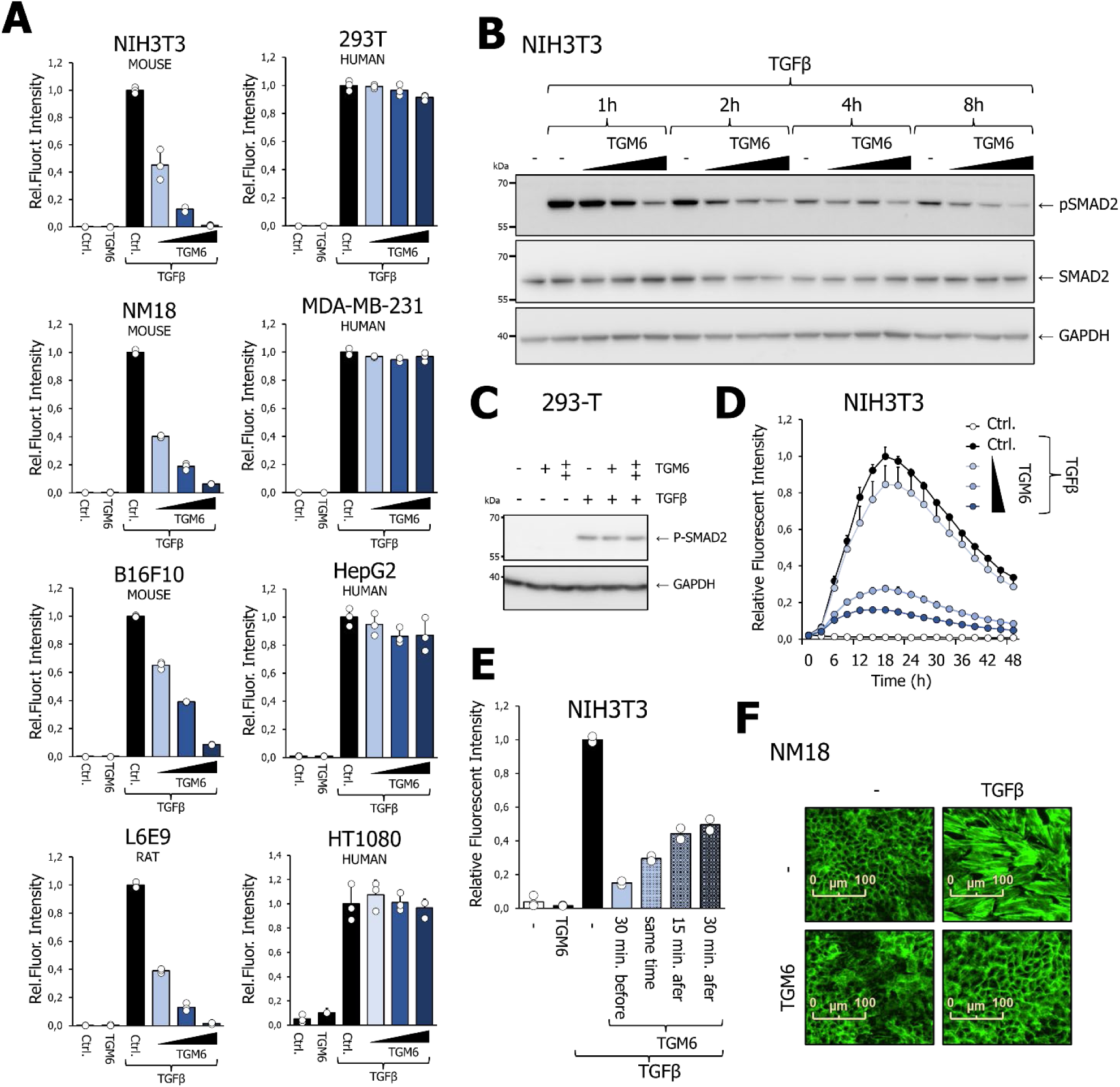
TGM6 potently antagonizes TGFβ/SMAD signaling in mouse, rat, but not human cells. (**A**) Effect of TGM6 on TGFβ/SMAD3-induced transcriptional response in mouse, rat, and human cells. Mouse NIH3T3 fibroblasts, NMuMG breast cells (NM18 clone), B16F10 melanoma, rat L6E9 myoblasts and human 293T epithelial, MDA-MD-231 breast cancer, HepG2 hepatoma and HT1080 fibrosarcoma cells that were engineered to express a CAGA-dynGFP reporter were pre-incubated for 30 min with different doses of TGM6 (10, 25, or 100 ng/ml) and were subsequently stimulated with 1 ng/ml TGFβ for 24h. The IncuCyte Life Cell Analysis System measured the GFP signal. (**B, C**) Effect of TGM6 on TGFβ-induced SMAD2 phosphorylation in NIH3T3 cells and HEK293T cells. (**B**) NIH3T3 cells were pre-incubated for 30 min with TGM6 (10, 25, or 100 ng/ml) and subsequently stimulated with 1 ng/ml TGFβ for 1, 2, 4 or 8h. (**C**) HEK293T cells were pre-incubated with 100 ng/ml TGM6 and thereafter treated with 1 ng/ml TGFβ. Cells were also treated with TGFβ alone. (**D**) Kinetic effect of TGM6 on NIH3T3 CAGA-dynGFP reporter activity; cells were treated with 1 ng/ml TGFβ and different doses of TGM6 (10, 25, or 100 ng/ml), and GFP signal was measured by IncuCyte Life Cell Analysis System. (**E**) Effect of different exposure time to TGM6 on inhibitory effect on TGFβ/SMAD signaling, NIH3T3 cells containing the CAGA-dynGFP reporter were pre-treated, treated at the same time, or with 100 ng/ml TGM6 and 1 ng/ml TGFβ (overnight). GFP reporter activity, as measured by the IncuCyte life analysis system after 21h of stimulation, is shown. (**F**) Effect of TGM6 100 ng/ml on TGFβ (1 ng/ml)-induced EMT in mouse NM18 breast cells.

### 2.2. Three amino acid differences between the mouse and human TGFBR2 extracellular domains are major determinants for TGM6 species specificity

TGM6 binds TGFBR2 with high affinity (*19*). To investigate the functional importance of this interaction, we mutated three TGM6 amino acid residues, i.e. Arg38, Ile78, and Tyr93 to Ala residues in TGM6 (TGM-6-mut), which are critical for TGFBR2 interaction based upon TGM6:hTGFBR2 extracellular domain (ECD) crystal structure [19], and analysed the effect of TGM-6-mut on TGFβ/SMAD signaling in mouse MFB-F11 fibroblasts. TGM6-mut was fully inactive, suggesting that TGM6-TGFBR2 interaction is essential for the antagonistic function of TGM6. When we compared the mouse (m) and human (h) TGFBR2 ECDs, we found 20/122 amino acid differences in the extracellular ligand binding domain, including 3 in the previously identified interface between TGM6-D3 and hTGFBR2 (Figure 2B). We therefore examined whether this dissimilarity is a crucial determinant of differential TGM6 activity. We ectopically expressed mTGFBR2 and hTGFBR2 (in a doxycycline-inducible manner) in NIH3T3 cells, which were deficient for mTGFBR2 due to CRISPR-CAS9 gene editing, and subjected the cells to TGFβ in the absence or presence of TGM6 and analyzed phosphorylated SMAD2 levels. We found that mTGFBR2, but not hTGFBR2, restored TGM6 responsiveness (Figure 2C), suggesting that mTGFBR2 is a critical mediator of TGM6 antagonist activity. We also noted that TGM6 weakly reduced TGFBR2 levels in mice but not human-expressing TGFBR2 NIH 3T3 cells (Figure 2C). Whether this decrease in TGFBR2 is part of the TGM6-induced antagonism of TGFβ signaling is addressed in subsequent experiments.

**Figure 2.**
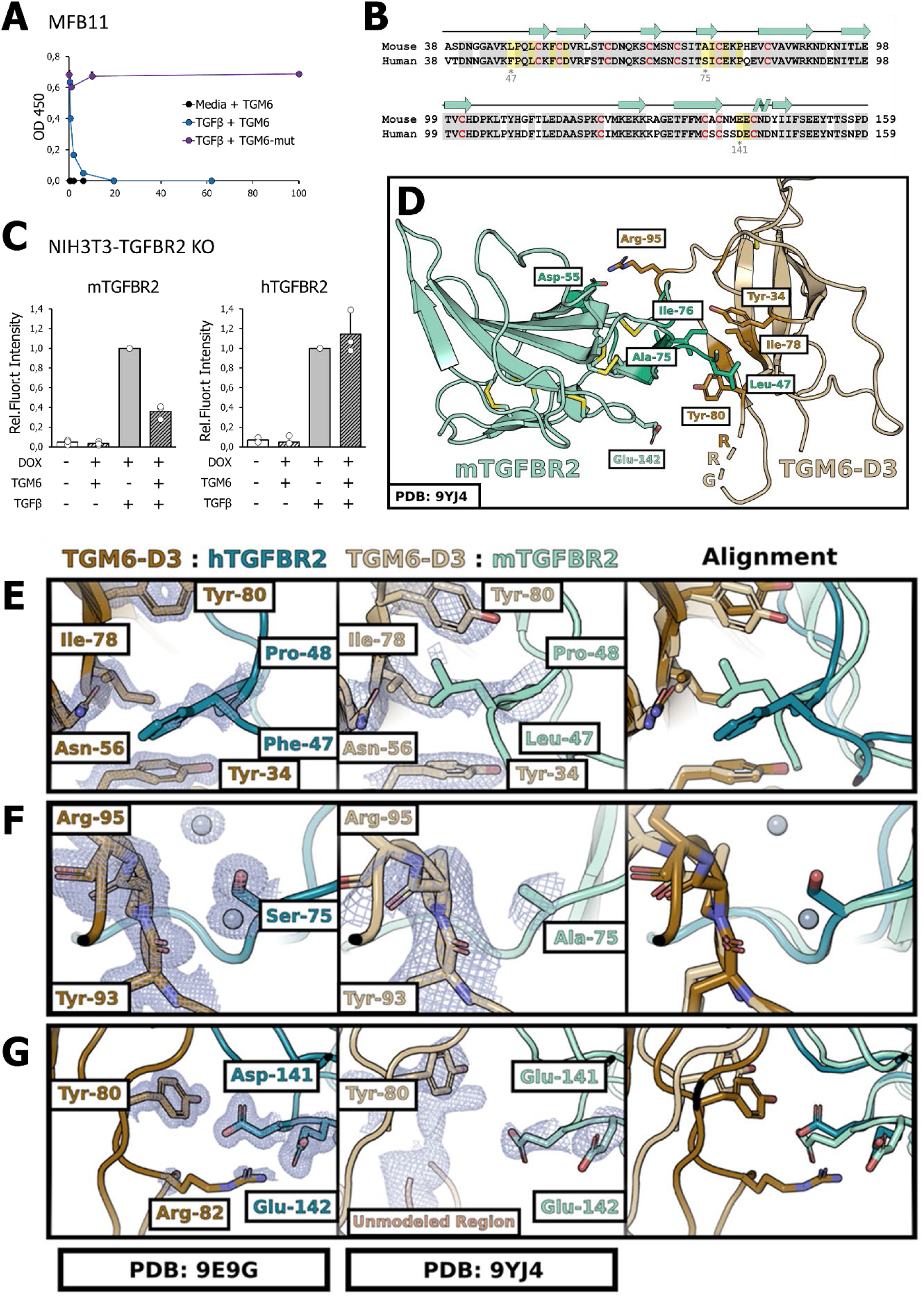
Three amino acid differences in the mouse and human TGFBR2 extracellular domain are responsible for TGM6 species specificity. (**A**) Effect of TGM6 (wildtype) or TGM6-mut (defective in TGFBR2 interaction) on TGFβ-induced activation of SMAD3 transcriptional activity using MFB-F11 reporter cells. (**B**) Amino acid sequence alignment of extracellular domains of mTGFBR2 and hTGFBR2. Identical non-interfacial and interfacial residues between mouse and human are highlighted with grey or dark yellow shading, respectively. Non-identical residues are not shaded. The three residues at the interface that are expected to differ between the complexes with hTGFBR2 (Phe47, Ser75, Asp141) and mTGRBR2 (Leu47, Ala 5, Glu141) are highlighted in light yellow with an asterisk. Conserved cysteine residues are indicated in red, and the β strands and α−helix are indicated on top of the primary amino acid sequence with blue arrows and blue helix, respectively. (**C**) Mouse, but not human, TGFBR2 can restore TGM6-induced antagonism of TGFβ/SMAD signaling in TGFBR2-knockout cells. NIH3T3 TGFBR2 knock-out cells were transduced with lentiviruses containing either mouse or human TGFBR2 under a doxycycline-inducible promoter. TGFBR2 expression was induced overnight by doxycycline, and cells were pretreated with 100 ng/ml TGM6 before stimulation with 1 ng/ml TGFβ; samples were analyzed by Western blotting for SMAD2 phosphorylation. The integrated results from three independent experiments are shown. (D) Overall structure of the mTGFBR2:TGM6-D3 ECD complex determined by X-ray crystallography at a resolution of 2.52 Å. mTGFBR2 ECD and TGM6-D3 are shaded teal and light brown, respectively. Loops that are not modeled due to weak density are indicated by dashed lines. Side chains of key interfacial residues, including those that differ relative to those in hTGFBR2, are shown, as are the intramolecular disulfide bonds (**E-G**). Comparison of TGM6-D3 in complex with human or mouse TGFBR2 at comparable interface regions at positions that differ between human and mouse TGFBR2. Left and middle panels show the model with density for the human and mouse complexes, respectively, while the right panel shows an alignment of the human and mouse models. In (**E**), Leu47 in mTGFBR2 is shown to fill the pocket formed by TGM6-D3 Tyr34, Ile78, and Tyr80 – in hTGFBR2, Phe47 in is unable to rotate inward to fill the pocket – hence this interface difference likely contributes to the preferential binding of mTGFBR2 to TGM6-D3. In (**F**), Ala75 in mTGFBR2 is shown to adopt essentially the same position as Ser^75^ in hTGFBR2, and so may only make a minor contribution to the preferential binding of mTGFBR2 to TGM6-D3. In (**G**), both Glu142 in mTGFBR2 and Asp142 in hTGFBR2 are positioned close to a highly mobile loop in TGM6-D3, and this interface difference might also only contribute to the preferential binding of mTGFBR2 to TGM6-D3 in a minor way.

To further investigate whether the increased antagonistic activity of TGM6 in mouse cells is due to preferential binding to mTGFBR2, we measured the binding affinity of purified TGM6-D3 for purified mTGFBR2 and hTGFBR2 using isothermal titration calorimetry (ITC) (Figure S1). These binding measurements, which were performed with duplicate titrations and global analysis of the integrated heats *(23,24),* showed that TGM6-D3 bound mTGFBR2 with 40-fold greater affinity compared to hTGFBR2 (15 nM vs. 604 nM, Table 1).

**Table 1.**
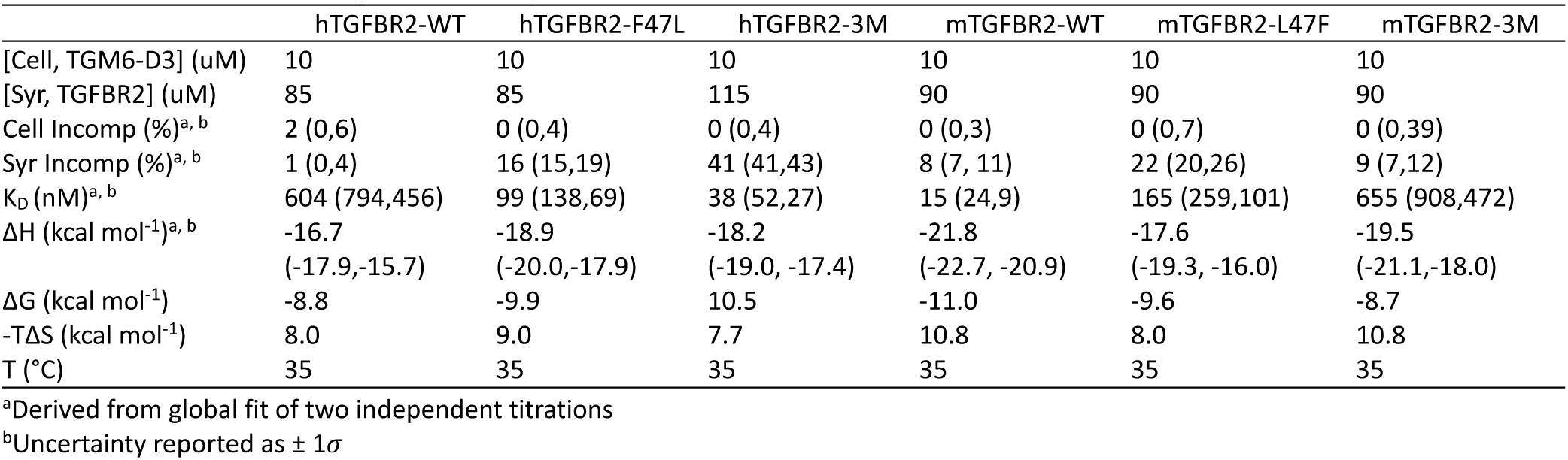
TGM6:TGFBR2 binding as assessed by IT.

The structure of the TGM6-D3:hTGFBR2 ECD complex [19] shows that only three interface residues are expected to differ between the complex with hTGFBR2 (Phe47, Ser75, Asp141) and mTGRBR2 (Leu47, Ala75, Glu141). Thus, we hypothesized that these three residues are responsible for the strong preferential binding of TGM6-D3 to mTGFBR2. To investigate this, we crystallized the TGM6-D3:mTGFBR2 complex and determined its structure at 2.52 Å resolution by X-ray crystallography (Table S1). This showed that the overall structure of the TGM6-D3:mTGFBR2 complex was very similar to the previously reported structure of the TGM6-D3:hTGFBR2 complex, with an overall root mean square deviation (RMSD) over all heavy atoms of 0.49 – 0.58 Å (Figure 2D, S2). The close similarity was unsurprising, given that mouse and human TGFBR2 are 83% identical across the entire ECD.

Through closer inspection of the structures of TGM6-D3 in complex with mTGFBR2 and hTGFBR2, we found that the side chains of the conserved interface residues engage TGM6-D3 in an essentially identical manner. There were nonetheless significant differences for one of the non-conserved interface residues, residue 47 (Figure 2E). In the structure of the mouse complex, Leu47 adopts a sidechain rotamer conformation that allows it to fill a hydrophobic pocket formed by TGM6-D3 Tyr34, Ile78, and Tyr80. In the human complex, Phe47 could enter the pocket only in a high-energy conformation due to its reduced conformational flexibility relative to leucine. In contrast to residue 47, only minor structural differences and no apparent changes in interactions are observed for residues 75 and 141 that might account for the preferential binding of TGM6-D3 for mTGFBR2 (Figure 2F, G).

To test our hypothesis that the preferential binding of TGM6-D3 for mTGFBR2 is engendered primarily by Leu47 (in place of Phe47 in hTGFBR2), we expressed and purified the hTGFBR2 F47L and mTGFBR2 L47F single amino acid variants and hTGFBR2 F47L, S75A, D141E and mTGFBR2 L47F, A75S, and E141D triple amino acid variants (hTGFBR2-3M and mTGFBR2-3M, respectively) and measured their binding affinity for TGM6-D3 using ITC. These binding measurements, which were performed with duplicate titrations and global analysis of the integrated heats *(23,24),* showed that the three residues were indeed responsible for the preferential binding, with the hTGFBR2-3M gain-of-function and the mTGFBR2-3M loss-of-function variants binding comparably to mTGFBR2-WT and hTGFBR2-WT, respectively (Table 1). The data further showed that residue 47 is the primary determinant of preferential binding, as the hTGFBR2-F47L and mTGFBR2-L47F variants gained and lost nearly two-thirds of the binding affinity relative to mTGFBR2-WT and hTGFBR2-WT, respectively (Table 1). Together, this data shows that TGM6 preferentially antagonizes signaling in mouse cells due to a 30-40 fold preference for binding mTGFBR2 over hTGFBR2.

### 2.3. TGM6 binds to TGFBR2 and Betaglycan and competes with TGFβ for TGFBR2 binding

In a search for TGM6 (co)receptors, we performed affinity crosslinking of iodinated TGM6 to cell surface proteins on responsive NIH3T3 cells (Figure 3A). Analysis of cell lysates immunoprecipitated with antibodies to TGFβ family type I (i.e., activin receptor-like kinase (ALK)1, –2, –3, –4, –5 (TGFBR1), type 2 receptors (i.e., activin type II receptor (ACTR2)A, ACTR2B, TGFBR2, bone morphogenetic protein type II receptor (BMPR2) and type III receptors (i.e endoglin and betaglycan)), revealed that as expected, only TGFBR2 from the type II receptors bound TGM6, but that none of the type I receptors or other type II receptors bound (Figure 3B). Interestingly, a betaglycan-TGM6, but not endoglin-TGM6, crosslinked complex was detected (Figure 3B). Importantly, in the TGM6 affinity-labeled cell lysate, besides the TGFBR2-TGM6 and betaglycan-TGM6 complex, an additional prominent TGM6 protein complex of about 600 kDa was present (Figure 3B). These results suggest that betaglycan and another unknown 600 kDa protein are co-receptors for TGM6 in NIH3T3 cells.

**Figure 3.**
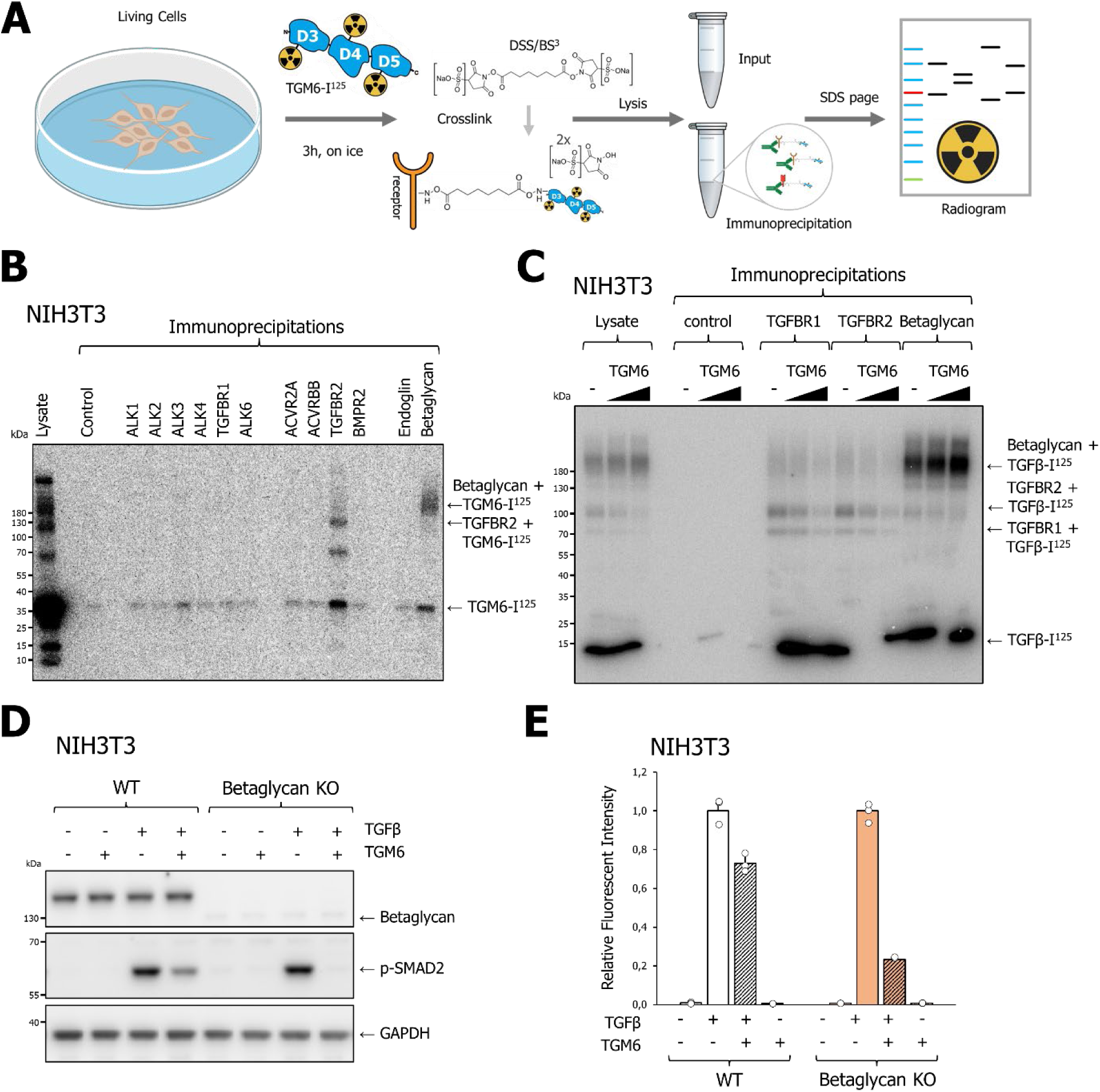
TGM6 competes with TGFβ for TGFBR2 interaction, and binds Betaglycan, a negative regulator of TGM6 function. (**A**) Schematic of the experimental flow of the affinity labeling experiment of NIH3T3 cells with iodinated TGM6. (**B**) Analysing the binding of TGM6 to TGFβ family type I, type II, and type III receptors. NM18 cells were incubated with iodinated TGM6, then crosslinked, lysed, and incubated overnight with receptor-specific antibodies. Thereafter, complexes were isolated using protein A beads, separated by SDS-PAGE, and detected by autoradiography. (**C**) Effect of TGM6 on radiolabeled TGFβ binding to TGFBR1, TGFBR2, and betaglycan. NIH3T3 cells were pre-incubated with 100 ng/ml TGM6 for 30 min before being incubated with iodinated TGFβ, then crosslinked, lysed, and incubated overnight with receptor-specific antibodies; finally, complexes were isolated using protein A beads, separated by SDS-PAGE and detected by autoradiography. (**D**) Western blot analysis for (lack of) betaglycan expression on CRISPR/CAS-generated betaglycan knock-out clones in NIH3T3 cells; two clones of each guide were analyzed for betaglycan expression. (**D**) Effect of betaglycan deficiency on the ability of TGM6 to antagonize TGFβ-induced CAGA-dynGFP reporter activity in NIH3T3 control clones, and betaglycan KO clones. Cells were pre-incubated with TGM6 (10 ng/ml) before stimulation with 1 ng/ml TGFβ for 21 h. (**E**) Effect of betaglycan deficiency on TGM6-mediated inhibition of TGFβ-induced SMAD2 phosphorylation in NIH3T3 control clones and betaglycan KO clones. Cells were pre-incubated with low-dose TGM6 (10 ng/ml) before stimulation with 1 ng/ml TGFβ for 1 h. Extended data for Figure 3D and E is shown in Figure S3C, S3D, and S3E.

We found that TGM6 binds with high affinity (15 nM) to mTGFBR2, and in the crystal structure of TGM6-D3-mTGFBR2 complex, the interface is remarkably similar to that of TGFβ-TGFBR2 (*19*) (Figure 2). Moreover, affinity binding assays with purified proteins revealed that TGM6 D3 competes with TGFβ for TGFBR2 binding [19]. We therefore investigated whether TGM6 can compete with TGFβ for mTGFBR on intact cells using affinity labeling of NIH3T3 cell surface proteins with iodinated TGFβ in the absence or presence of different amounts of TGM6. Cell lysates immunoprecipitated with TGFBR1, TGFBR2, or betaglycan-specific antibodies were analyzed. Of note, these affinity labeling experiments were conducted by keeping NIH3T3 cells on ice, thereby preventing TGFβ-induced TGFBR internalization. We observed that increasing amounts of TGM6 inhibited the TGFβ-mTGFBR1 and TGFβ-mTGFBR2, yet not TGFβ-betaglycan interactions (Figure 3C). TGM6 binds to TGFBR2, not TGFBR1; however, because TGFβ binding to TGFBR1 requires TGFBR2 (4), TGM6 also decreases TGFβ-mTGFBR1 interaction. The competition between TGM6 and TGFβ for TGFBR binding is consistent with the function of TGM6 as a TGFβ antagonist.

### 2.4. Betaglycan inhibits TGM6-induced antagonism of TGFβ signaling

To address the functional role of TGM6-betaglycan interaction, we derived replicate knock-out clones (1A, 1B, 2A, and 2B; 1 and 2 are corresponding to different guide RNAs, and A and B refer to different independent clones) for betaglycan in NIH3T3 cells using CRISPR-CAS9 gene editing (Figure S3A). When we analyzed the TGM6-induced repression of TGFβ-induced SMAD2 phosphorylation (Figure 3D, S3B) and TGFβ-induced SMAD3/4 transcriptional response and in control versus betaglycan knock-out clones (Figure 3E, S3C), we observed that deficiency of betaglycan greatly potentiated the TGM6-induced inhibitory effect on TGFβ signaling (Figure 3D, 3E, S3B, S3C). Thus, betaglycan is a negative regulator of TGM6.

Betaglycan comprises two major domains: an N-terminal Orphan domain and a C-terminal Zona Pellucida (ZP) domain [25]. To map which betaglycan domain(s) interact(s) with TGM6, we used rat (r) L6E9 myoblast cells deficient in betaglycan. L6E9 myoblasts are responsive to TGM6 (Figure 1A). We first confirmed that mbetaglycan and rbetaglycan are expressed after transfection with expression plasmids (Figure S3D) and bind TGM6 in L6E9 cells (Figure S3E). Ectopic expression of rBetaglycan wildtype or mutants lacking either the Orphan or ZP domains (Figure S3F) all interfered with the ability of TGM6 to exert its inhibitory effect on TGFβ-induced SMAD3/4 transcriptional response (Figure S3G) and TGFβ-induced SMAD2 phosphorylation (Figure S3H). Thus, both orphan and ZP domains are important for TGM6-Betaglycan interplay.

### 2.5. Identification of low-density lipoprotein receptor-related protein (LRP)1 as TGM6 co-receptor, which is a key determinant for TGM6-induced antagonism of TGFβ signaling

To identify cell-surface binding proteins in an unbiased manner, biotinylated TGM6 was added to NIH3T3 cells, and the cell lysates were subjected to a pulldown assay using neutravidin beads. The proteins bound were trypsinized, and peptide fragments analyzed by mass spectrometry (Figure 4A). As expected, TGFBR2, known to interact with TGM6, was identified as a highly abundant protein in the pulldown (Figure 4B). Among the other hits, low-density lipoprotein receptor-related protein 1 (LRP1) caught our attention (Figure 4B), as it is a large 600 kDa type I receptor glycoprotein involved in receptor-mediated endocytosis [17]. Like other members of the LDLR protein family, LRP1 contains cysteine-rich complement-type repeats, epidermal growth factor (EGF) repeats, and β propeller domains, a transmembrane domain, and an intracellular domain. LRP1 contains four ligand-binding domains numbered I to IV and multiple cysteine-rich complement-type repeats. LRP1 is processed by furin, thereby producing two non-covalently bound chains. i.e. a 515 kDa α-chain and an 85 kDa β-chain [26]. Notably, the 600 kDa TGM6-protein complex that was detected in the cell lysate (Figure 3B) corresponds to the expected size of TGM6 cross-linked with (heavily glycosylated) LRP1α (and LRP1β) (Figure 3B).

**Figure 4.**
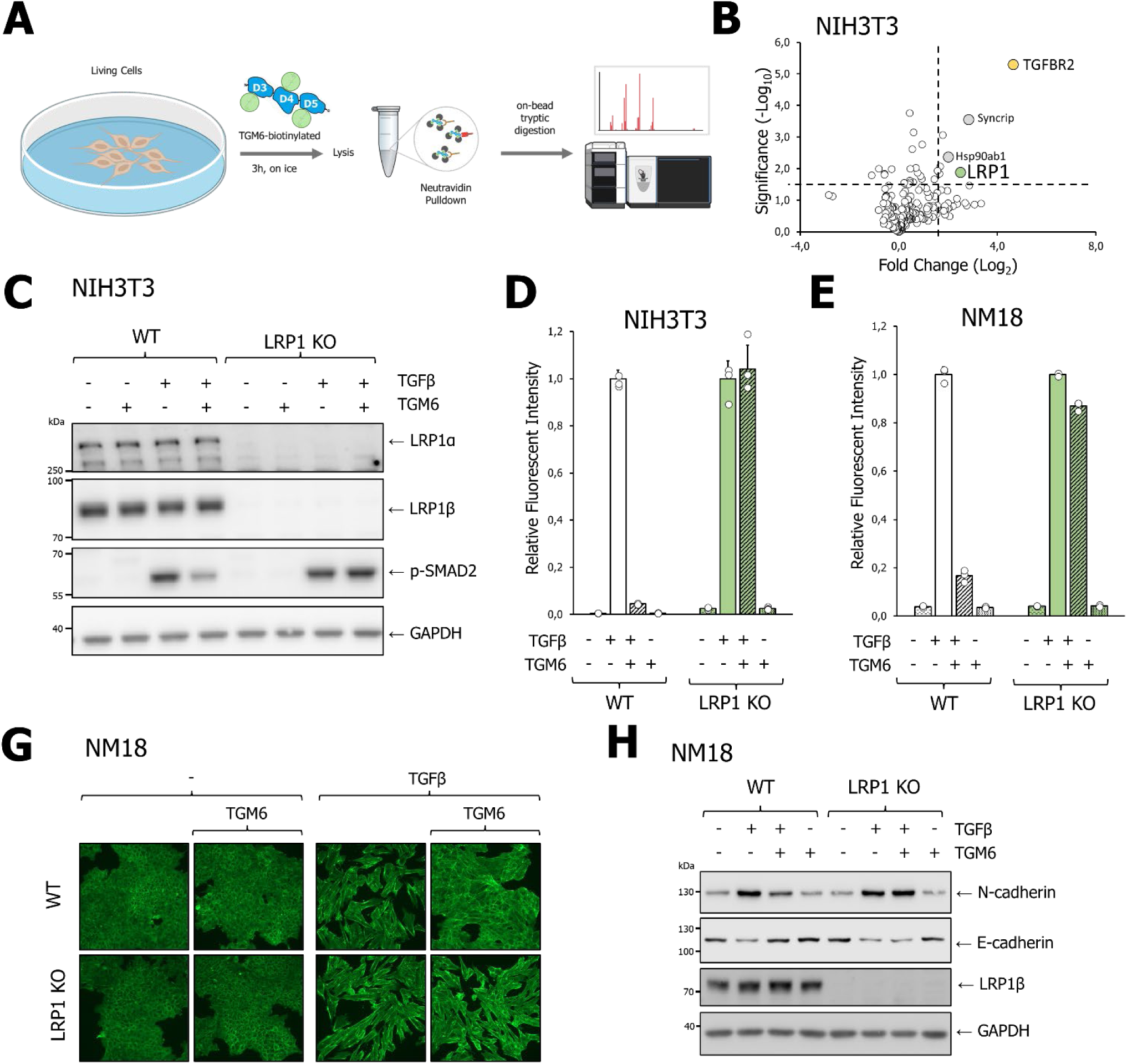
Identification of LRP1 as TGM6 co-receptor. (**B**) Schematic representation of the experimental flow for the identification of TGM6 interactors using biotinylated TGM6. NIH3T3 cells were incubated with biotinylated TGM6, cells were lysed, and TGM6-biotin and interaction partners were precipitated using neutravidin beads and analysed by mass spectrometry. (**B**) Analysis of TGM6 binding partners in NIH3T3 cells by mass spectrometry. Fold change is calculated relative to control cells undergoing the same procedure without TGM6-biotin addition. TGM6 interaction partners TGFBR2 and LRP1 are highlighted with yellow and blue circles. (**B and C**) Western blot analysis for (lack of) LRP1 expression in CRISPR/CAS-generated NIH3T3 (**B**) and NM18 (**C**) LRP1 knock-out clones; 2 clones of each guide were analyzed for LRP1 expression. (**C, D,** and **ED**) Effect of LRP1 deficiency in NIH3T3 cells (C and D) and NM18 cells € on antagonism of TGM6 on TGFβ-induced SMAD2 phosphorylation (C) and TGFβ/SMAD3 transcriptional activity in NIH3T3 cells (D) and NM18 cells (E). Control clones and LRP1 KO clones were pre-incubated with TGM6 (100 ng/ml) for 30 min before stimulation with 1 ng/ml TGFβ for 1 hour (for Western blot assay) and 21 hours (for transcriptional reporter assay). Extended (supporting) data for Figure 4 C, D, and E are shown in Figure S4A, S4B, S4C, and S4D. (**G**) Effect of LRP1 deficiency on antagonism of TGM6 on TGFβ-induced morphological transition and actin stress fiber formation in NM18 cells. NM18 cells were pre-treated with 100 ng/ml TGM6 for 30 min and subsequently stimulated with 1 ng/ml TGFβ, and after 2 days, cells were fixed and stained with Phalloidin-ALEXA-488. (**H**) Effect of LRP1 deficiency on TGFβ (1 ng/ml)-induced EMT (after 2 days of stimulation) as measured by Western blot analysis of epithelial E-Cadherin and mesenchymal N-Cadherin expression.

Similar to our strategy for betaglycan, we investigated the functional role of TGM6-LRP1 interaction by knocking out LRP1 in NIH3T3 cells and NM18 cells using CRISPR-CAS9 gene editing. We again generated 4 LRP1 knockout clones (KO1A, KO1B, KO2A, and KO2B) in NIH3T3 cells (Figure S4A) and in knockout LRP1 pools (KO1 and KO2) of NM18 cells (Figure S4B). KO1A and KO1B were generated using different guides, and A and B refer to independent clones/pools. TGM6 was found to lose all inhibitory effects on the TGFβ-induced pSMAD2 phosphorylation in LRP1-KO cells (Figure 4C, S4C), a result confirmed by the intact TGFβ/SMAD3 transcriptional response in LRP1-KO cells in the presence of TGM6 (Figure 4D, S4D). Similar results were obtained using NMuMG breast epithelial cells, clone NM18 (Figure 4E and S4E). NMuMG (NM18) cells are a frequently used model system for examining TGF-β-induced EMT (18). The TGFβ-induced morphological change and formation of actin stress fibers observed in control cells were blocked by TGM6 in control/wild-type cells, but not in LRP1-deficient NM18 cells (Figure 4G). Consistently, whereas TGM6 blocked the downregulation of the epithelial marker E-cadherin and the upregulation of the mesenchymal marker N-cadherin in control/wild-type cells, this was not observed in LRP1 knock-out cells (Figure 4H). Thus, LRP1 is a critical determinant of the cell-type-specific potency of TGM6, presumably by increasing its cellular avidity relative to cell types that lack LRP1 expression.

### 2.6. TGM6 interacts via separate domains with LRP1 and TGFBR2, and TGM6-betglycan requires TGFBR2

To further validate the TGM6 interaction with LRP1 and investigate the possible interplay by which the three different receptors and co-receptors interact with TGM6, we compared the binding of TGM6 to cell surface proteins in wild-type NIH3T3 cells or NIH3T3 cells deficient in LRP1, betaglycan, or TGFBR2 (Figure 5A-C). Affinity labeling with iodinated TGM6 in control/wild-type cells, and immunoprecipitation of cell lysates with an LRP1 antibody, confirmed that TGM6 binds LRP1. Immunoprecipitation for TGFBR2 or betaglycan in cell lysates of LRP1 knock-out cells that were affinity labeled with iodinated TGM6 revealed that TGM6, in the absence of LRP1, is still able to interact with TGFBR2 and betaglycan, albeit weaker than in control/wild-type cells (Figure 5D). We observed no co-immunoprecipitation, indicative of heteromeric complex formation, of TGFBR2 or betaglycan with LRP1 (Figure 5D). Similar affinity labeling followed by receptor immunoprecipitation experiments performed with iodinated TGM6 on NIH3T3 betaglycan knock-out cells or control/wild-type cells showed that TGM6 binding to TGFBR2 is not different in knock-out versus control/wild-type cells, while TGM6 binding to LRP1 is slightly increased in betaglycan-deficient cells (Figure 5E). Affinity labeling of cell surface receptors with iodinated TGM6 on NIH3T3 cells deficient in TGFBR2 versus control/wildtype revealed that betaglycan binding is nearly absent and TGM6-LRP1 interaction is unperturbed (Figure 5F). Thus, TGM6 binds TGFBR2 and LRP1 without needing betaglycan, but betaglycan requires TGFBR2 to interact with TGM6.

**Figure 5.**
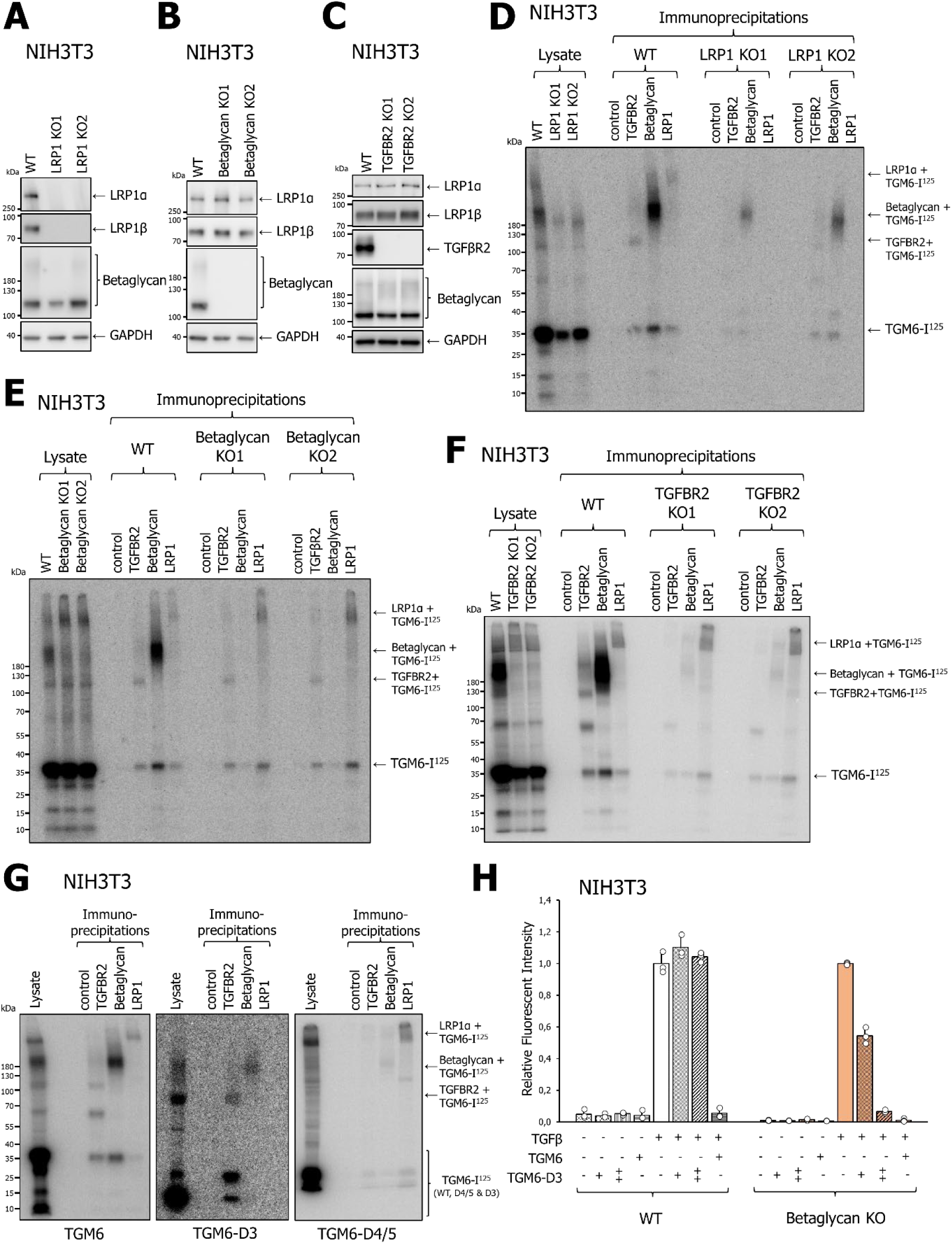
TGM6 D4/5-LRP1 interaction occurs independently of TGBR2 and Betaglycan, but the interaction of TGM D3-Betaglycan requires TGFBR2. (**A-C**) Western blot analysis of LRP1, betaglycan, and TGFBR2 expression in NIH3T3 LRP1 (A), NIH3T3 betaglycan (B), or NIH3T3 TGFBR2 (C) knock-out clones (1 and 2). (**D-F**) Effect of LRP1 deficiency (**D**), betaglycan deficiency (**E**), or TGFBR2 deficiency (**F**) on TGM6 binding to (co)receptors. Iodinated TGM6 was cross-linked to NIH3T3 control cells or various KO cells, and cell lysates were immunoprecipitated with the indicated antibodies. Total cell lysates were analysed. Signals were analysed by autoradiography. (**G**) Analysis of wild-type TGM6, TGM6-D3, and TGM6-D4/4 to (co)receptors. Iodinated TGM6 (left panel), TGM6-D3 (middle panel), or TGM6-D4/5 (right panel) were used to affinity label cell surface proteins on NIH3T3 cells, and cell lysates were immunoprecipitated using the indicated antibodies. Total cell lysates were also analysed. Signals were analysed by autoradiography. (**H**). Effect of betaglycan deficiency on the ability of TGM6-D3 to antagonize TGFβ-induced CAGA-dynGFP reporter activity in NIH3T3 control clones, and betaglycan NIH3T3 KO clones. Cells were pre-incubated with TGM6 (100 ng/ml) or TGM6-D3 (1000 or 5000 ng/ml) for 30 min before stimulation with 1 ng/ml TGFβ for 21 h. Extended data for Figure 5H is shown in Figure S5.

To further consolidate and expand these findings, we compared the binding of intact TGM6, TGM6-D3, or TGM6-D4/5 truncations to the three (co)receptors (Figure 5G). As expected, intact TGM6 interacts with TGFBR2, betaglycan, and LRP1. TGM6-D3 weakly interacts with TGFBR2 and betaglycan (and not LRP1), and TGM6-D4/5 interacts with LRP1 (and not TGFBR2 and betaglycan) (Figure 5G). Thus, TGM6-D3 is both required and sufficient for interaction with TGFBR2 and betaglycan, while TGM6-D4/5 is essential and sufficient for LRP1 binding.

We previously demonstrated that TGM6-D3 is insufficient to inhibit TGF-β signaling in fibroblasts *(19)*. As betaglycan inhibits TGM6 function, we compared the effect of TGM6-D3 in wild-type versus betaglycan knock-out cells to investigate if, under these conditions, TGM6-D3 can exert TGFβ antagonism. TGM6-D3 inhibited the TGFβ-induced SMAD3 transcriptional response in a dose-dependent manner, albeit with very high doses of TGM6 (1000 to 5000 ng/ml) in NIH3T3 cells depleted of betaglycan, but not in wild-type NIH3T3 cells (Figure 5H, S5).

### 2.7. TGM6 D4/5 interacts with LRP1 via its LDLaIV cluster

To map the LRP1 domain that interacts with TGM6 D4/5, we generated a series of mLRP1 deletion constructs. All mLRP1 deletion constructs retained a signal peptide and a transmembrane domain to enable cell surface expression (Figure 6A). Expression of LRP1 constructs after transfection of 293T cells was confirmed by Western Blot analysis of cell lysates (Figure 6B). Affinity labeling of LRP1 (deletion) expression constructs with iodinated TGM6 revealed that TGM6 binds to LRP1-D4 (Figure 6C, image on the left). This region contains the LDLaIV cluster, of which the tandem repeats of the cysteine-rich lipoprotein receptor class A (LDLa) can form binding sites for ligands [26]. TGM6 was found to interact with the LDLaIV cluster, but not with a related LDLaII cluster used as a specificity control (Figure 6C, image on the right). Ectopic expression of hLRP1 or mLRP1 D4 in LRP1-deficient NIH3T3 cells rescued the ability of TGM6 to inhibit TGFβ signaling as measured by SMAD3-dependent transcriptional response (Figure 6D) and TGFβ-induced SMAD2 phosphorylation (Figure 6E). Ectopic expression of hLRP1 D4 was as potent as mLRP1 D4 to rescue the TGM6 inhibitory effect on TGFβ signaling in NIH3T3 LRP1 KO cells (Figure 6D). Furthermore, we found that mLRP1-LDLaIV (with the transmembrane (TM) domain) rescued the TGM6 inhibitory effect on TGFβ/SMAD signaling, albeit less efficiently than mLRP1-D4 (Figure 6E). Next, we investigated the impact of soluble mouse or human LRP1-LDLaIV on TGM6-induced inhibition of TGFβ/SMAD signaling. Both soluble proteins were equally expressed in the conditioned media of transfected HEK293T cells (Figure S6A). We found that soluble hLRP1-LDLaIV or mLRP1-LDLaIV inhibits TGM6 inhibitory effect on TGFβ/SMAD3 transcriptional response in a dose-dependent manner (Figure 6F and S6B). LRP1-LDLaIV acts as a trap that sequesters TGM6, preventing its interaction with LRP1. Our results support the notion that the differential binding of TGM6 to mTGFBR2 versus hTGFBR2 (Figure 2) and not mLRP1 versus hLRP1, is the critical determinant for mouse versus human cell selectivity of TGM6 responses. Consistent with our biochemical results, Alphafold3 predicted with high confidence specific interactions between TGM6-D3 and TGM6 D4/5 with mLRP1-LDLa IV and mTGFBR2, respectively (Figure 6G). Taken together, our results indicate that the LDLaIV cluster is both sufficient and required for TGM6 interaction, and that TGM6-D4/5 interaction with LRP1 occurs independently of TGM6-D3.

**Figure 6.**
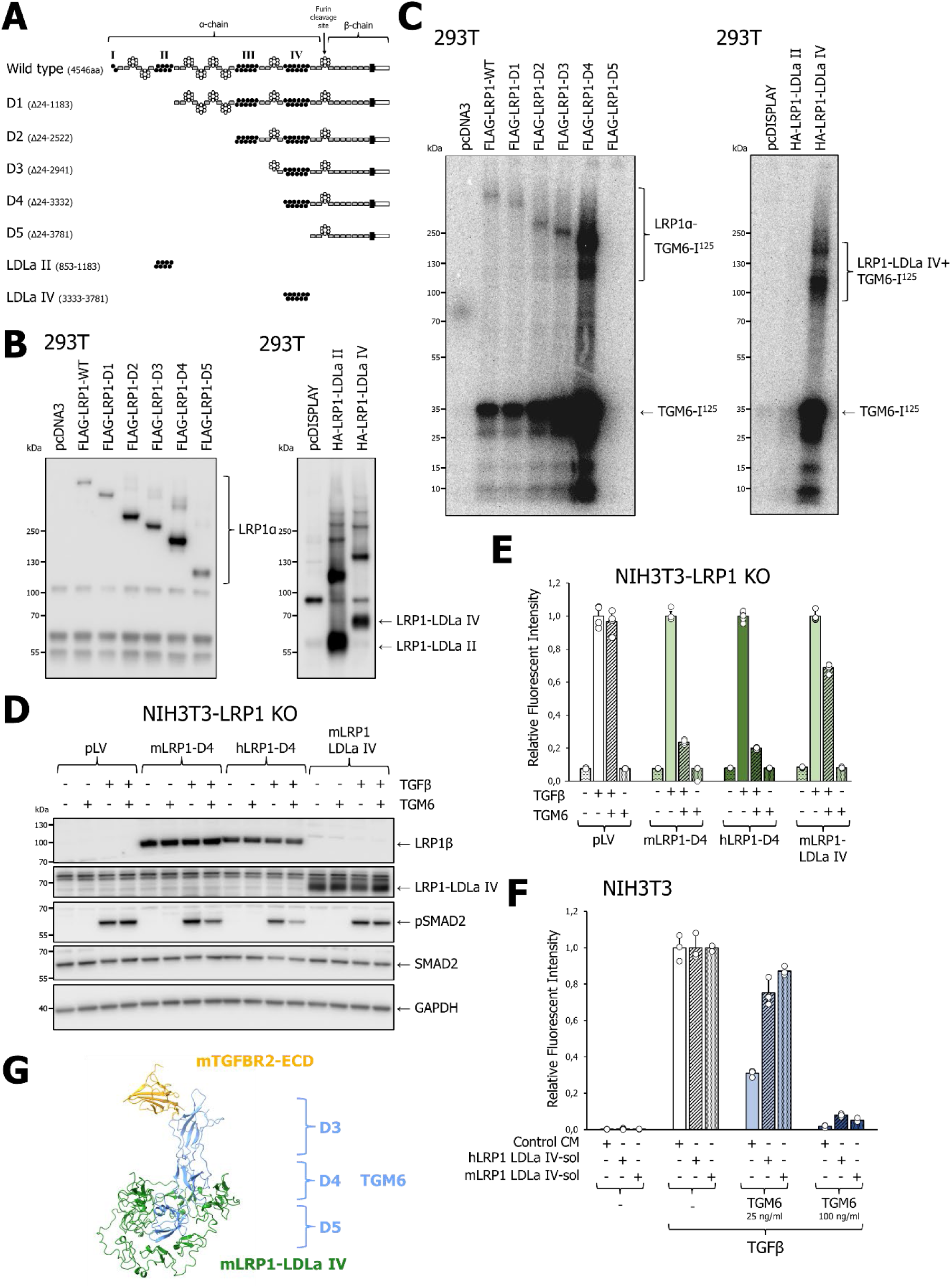
LRP1 LDLaIV cluster interacts with TGM6-D4/5. (**A**) Schematic of the LRP1 constructs, WT, D1, D2, D3, D4, and D5 (pcDNA3), LDLaII, and LDLaIV (pDISPLAY). This figure was modified from [65]. (**B**) Expression analysis of LRP1 wild-type and deletion constructs by Western blot analysis of proteins that were analysed by immunoprecipitation in (C). (**C**) Analysis of TGM6 binding to LRP1 deletion constructs. HEK293T cells overexpressing the different constructs were used to cross-link with radio-labeled TGM6. LRP1 was immunoprecipitated using anti-FLAG (deletion constructs on the left) or anti-HA (LDLa clusters on the right). (**D** and **E**) Effect of hLRP1-D4 and mLRP1-D4 on TGM6 (100 ng/ml)-induced inhibition of TGFβ (1 ng/ml) signaling in LRP1 KO cells as measured by CAGA-dynGFP reporter activation after 21 hours (**D**) and SMAD2 phosphorylation after 1 hour (**E**). NIH3T3 KO cells were transduced with lentiviruses containing the different LRP1 constructs. The cells were pre-incubated with TGM6 (100 ng/ml) before stimulation with 1 ng/ml TGFβ for 21 hours (D) or 1 hour (E). (**F**) Effect of soluble m LRP1-LDLaIV and hLRP1-LDLaIV on TGM6-induced inhibition of TGFβ-induced signaling. TGM6 was pre-incubated with conditioned media containing soluble LRP1 LDLaIV for 30 min. Subsequently, this was used to pre-incubate NIH3T3 cells for 30 min, followed by stimulation with 1 ng/ml TGFβ for 21 hours. (**G**) AlphaFold prediction of TGM6 in complex with mTGFBR2 ECD and mLRP1-LDLaIV, showing independent binding of TGM6-D3 to mTGFBR2 and TGM6-D4/5 to mLRP1-LDLaIV.

### 2.8. TGM6 acts *in-cis* and not *in-trans*

TGM6 has D3 and D4/5 domains interacting simultaneously but independently with TGFBR2 and LRP1, respectively. This allows them to engage target TGFBR2^+^LRP1^+^ cell types *in-cis* with high cellular selectivity and avidity. The modular design may also provide sufficient flexibility for TGM6 to interact with TGFBR2 and LRP1 in different cell types and act *in-trans*, thereby inhibiting TGFβ signaling in LRP^-^ cells that neighbor LRP1^+^ cells. To study if TGM6 acts *in-cis* and/or *in-trans*, we used NIH3T3-CAGA-dynGFP cells (TGFBR2^+^LRP1^+^) and NIH3T3-CAGA-mCHERRYd2 deficient in LRP1 (TGFBR2^+^LRP1^-^), either as mono– or mixed cultures in two different ratios (Figure S7A), and challenged them with TGM6 and/or TGFβ. As NIH3T3-CAGA-dynGFP and NIH3T3-CAGA-mCHERRYd2 express TGFBRs, TGFβ will induce CAGA-driven reporter activity in the two NIH3T3 cell types equally. Analysis of CAGA-dynGFP and CAGA-mCHERRYd2 levels revealed that, as expected, TGM6 potently inhibited TGFβ-induced CAGA-dynGFP response in the absence or presence of TGFBR2^+^LRP1^-^ cells, and TGM6 did not inhibit TGFβ-induced CAGA-mCHERRYd2 response in monoculture of TGFBR2^+^LRP1^-^ cells. CAGA-mCHERRYd2 signals were not inhibited when NIH3T3-CAGA-dynGFP and NIH3T3-CAGA-mCHERRYd2 were co-cultured (Figure S7B and C). To increase interactions between NIH3T3-CAGA-mCHERRYd2 and NIH3T3-CAGA-dynGFP, we used a five-fold excess of NIH3T3-CAGA-mCHERRYd2 over NIH3T3-CAGA-dynGFP. As CAGA-dynGFP is still potently decreased by TGM6 under these latter conditions, TGFBR2 in LRP1-deficient cells does not act as a sink to prevent TGM6 engagement with TGFBR2 on LRP1^+^ cells. These results suggest that TGM6 only acts *in-cis* and is effective on LRP1-expressing target cells, but not co-cultured LRP1^-^ TGFBR2^+^ cells (Figure S7B and C).

### 2.9. TGM6 antagonizes TGFβ signaling by inducing LRP1-dependent TGFBR2 lysosomal degradation

LRP1 is involved in receptor endocytosis and in lysosomal degradation [26]. We therefore set out to explore whether TGM6 engagement with TGFBR2 and LRP1 induces TGFBR2 degradation. Indeed, we noted that in response to TGM6, mTGFBR2 expression was reduced in NIH3T3 cells (Figure 2B). Challenging NIH3T3 and NM18 cells with TGM6 decreased TGFBR2, yet not TGFBR1 levels (Figure 7A). No effect of TGM6 on LRP1 expression was found. TGM6 stimulation decreased betaglycan expression in NIH3T3 cells but not NM18 cells (Figure 7A). The agonist TGFβ inhibited TGFBR2 in both cell types, and thus, the TGFBR2 downregulation is not per se associated with an antagonistic effect. TGFβ stimulated TGFBR1 levels in NM18 cells (Figure 7A). Next, we examined the impact of TGM6 and/or TGFβ on TGFBR2 levels in control and LRP1-deficient NIH3T3 cells. We observed that TGM6 and TGFβ inhibit TGFBR2 expression in control/wild-type cells (Figure 8B). In LRP1-deficient cells, TGM6 was unable to downregulate TGFBR2, but TGFβ remained proficient in TGFBR2 downregulation (Figure 7B). Consistent with these results, we observed that TGM6 inhibits SMAD2 phosphorylation, and that this effect is more pronounced if the TGFβ agonist is added after 360 min rather than 30 min in NH3T3 and NM18 cells (Figure 7C). While competition between TGFβ and TGM6 for TGFBR2 occurs immediately, the TGM6-induced TGFBR2 downregulation is more pronounced at 6 hours than at 1 hour of treatment.

**Figure 7.**
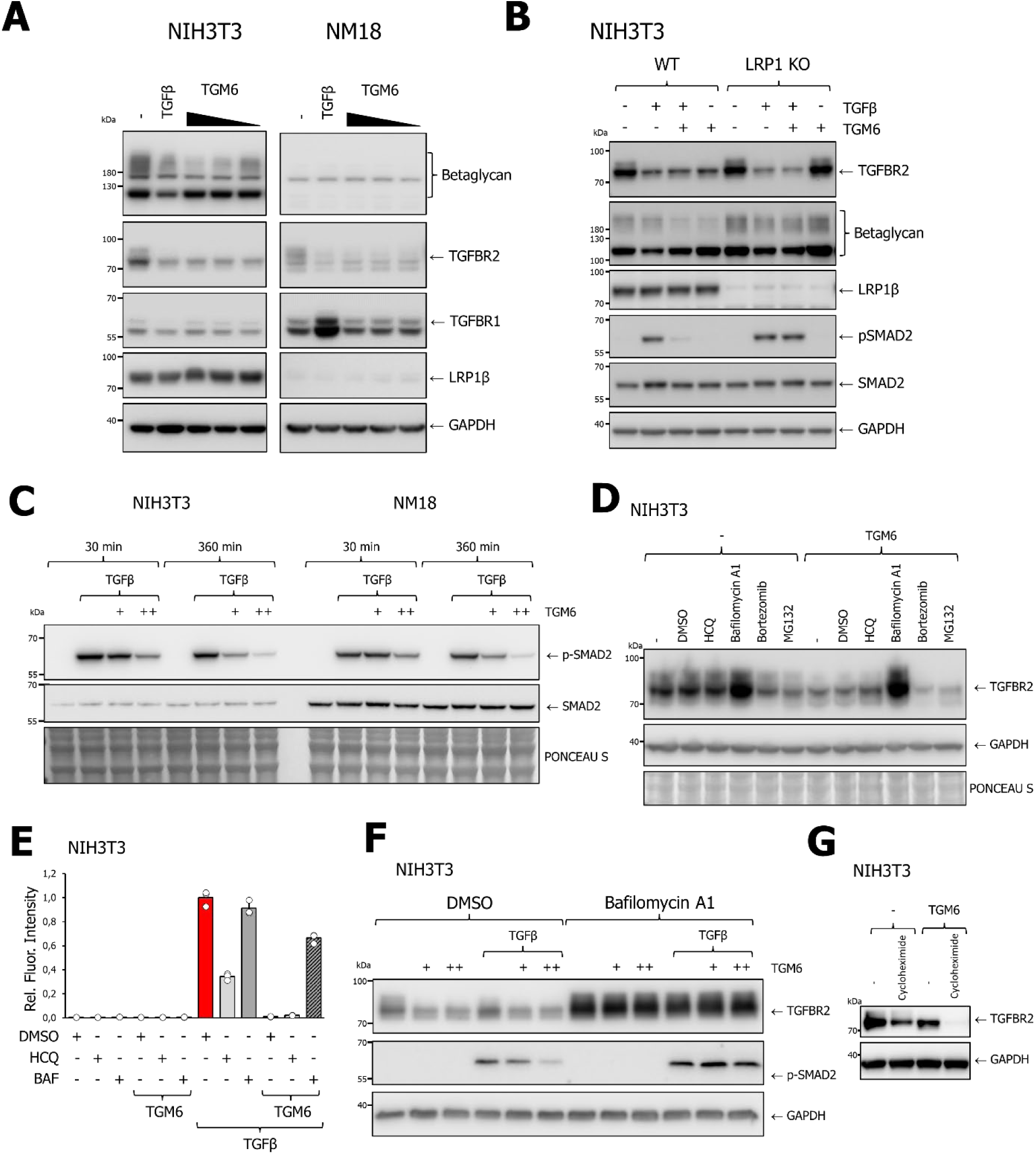
TGM6 antagonizes TGFβ signaling by inducing LRP1-dependent TGFBR2 lysosomal degradation. (**A**) Effect of TGM6 on expression of betaglycan, TGFBR1, LRP1β in NIH3T3 (Left) and NM18 cells (Right). Cells were treated with TGM6 (5, 50, or 500 ng/ml) or 1 ng/ml TGFβ for 6 hours, and cell lysates were thereafter analysed by Western blotting. (**B**) Effect of TGM6 (100 ng/ml) or TGFβ (1 ng/ml) on expression of betaglycan, TGFBR1, LRP1β, phosphorylated SMAD2 (pSMAD2), and SMAD2 expression levels in control /wild-type and LRP1 KO NIH3T3 cells. Cells were stimulated for 6 hours with the ligand. (A) and (B) were run on the same gel and analysed in the same manner (**C**) Effect of TGM6 on TGFβ-induced SMAD2 phosphorylation in NIH3T3 cells. 100 ng.ml TGM6 was added either 30 min or 360 min prior to the addition of 1 ng/ml TGFβ. (**D**) Effect of DMSO vehicle control, hydroxychloroquine (HCQ, 20 µM), bafilomycin A1 (20 nM), bortezomib (10 nM), and MG132 (5 µM) on TGM6-induced downregulation of TGFBR2 expression. Compounds were added 1 hour before TGM6 (100 ng/ml) for 6 hours. (**E**) Effect of TGM6 on TGFβ-induced SMAD3 transcriptional response in the presence of DMSO vehicle control, hydroxychloroquine (HCQ, 20 µM), or bafilomycin (20 nM). NIH3T3 cells were challenged with 100 ng/ml TGM6 (100 ng/ml) and/or TGFβ (1 ng/ml). The CAGA-dynGFP transcriptional response was measured after 20 hours. (**F**) Effect of inhibition of TGM6 (100 ng/ml) on TGFβ (1 ng/ml for 3 hours) induced SMAD2 phosphorylation in the absence or presence of bafilomycin A1 (20 nM, pre-treatment 1 hour). Pretreatment of TGM6 was 3 hours. (**G**) Effect of TGM6 (100 ng/ml) and/or cycloheximide (25 µg/ml) on TGFR2 expression, cells were pretreated with cycloheximide for 30 min before TGM6 (2 hours). Treatment with compounds was xx hours. (B-D, F, G) Expression analysis was performed as in (A).

**Figure 8.**
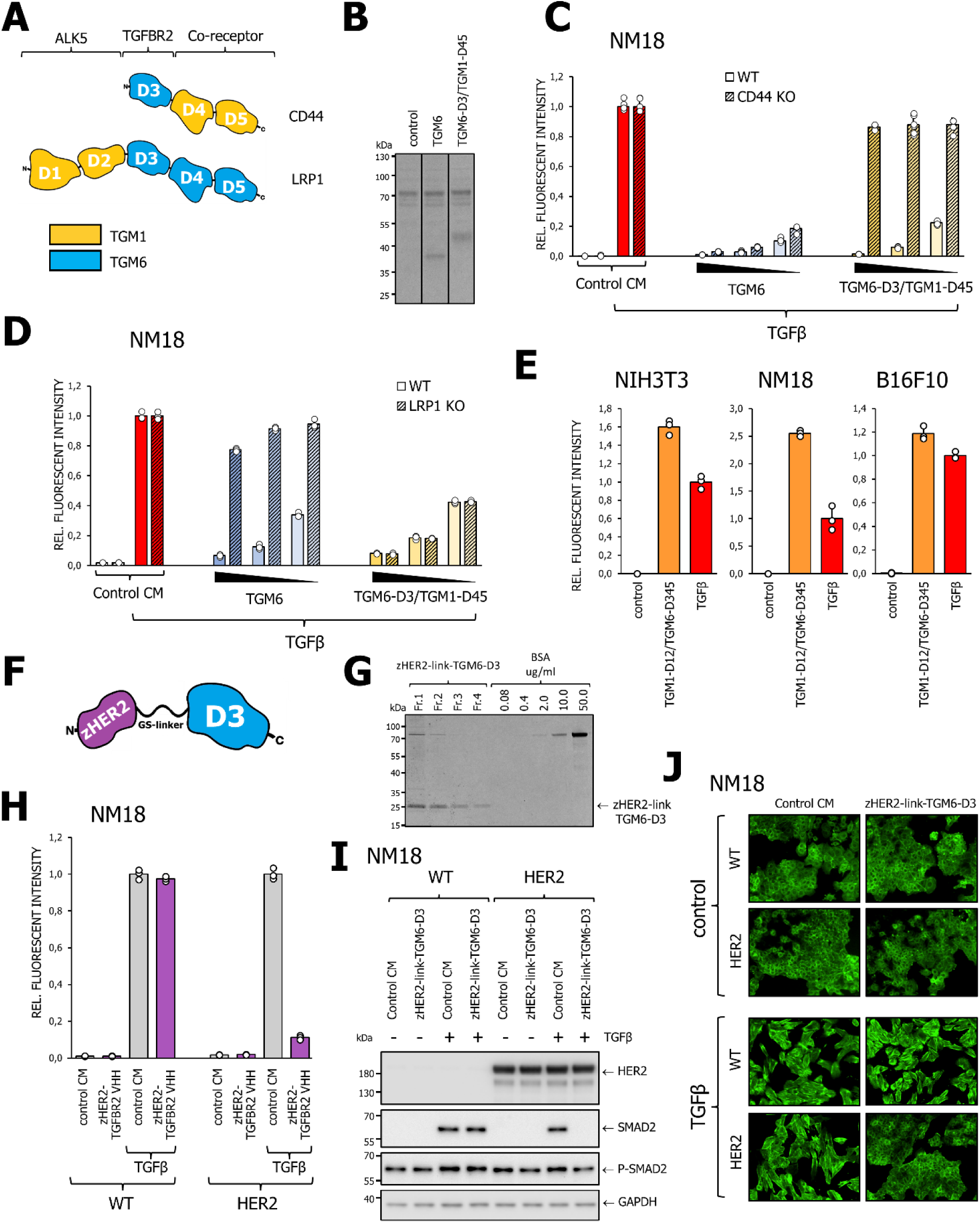
TGM6 chimeras change cell specificity and change TGM6 antagonist into a potent TGFβ agonist. (**A**) Schematic of the TGM1-TGM6 fusion proteins that were generated. (**B**) Coomassie stain of SDS polyacrylamide gel to analyze the expression and purity of TGM proteins and fusions in the conditioned media of transfected 293T cells. All three samples were run on the same gel. Lines indicate where the gel was cut. (**C**, **D**) Test whether the functionality of TGM6 can be switched by replacing its D4/5 domains with those of TGM1. Effect of TGM6 and TGM6-D3-TGM1-D45 chimera on the inhibitory potential of TGFβ signaling in (**C**) NM18 CAGA-dynGFP WT vs CD44 KO cells (single cell clone) and (**D**) NM18 CAGA-dynGFP WT vs LRP1 KO cells (pool). The GFP signal after 21 hours was measured by IncuCyte. (**E**) Analysis of fusing TGM6 to TGM1-D12 indicated that TGM6 can be converted from an antagonist into an agonist in NIH3T3, NM18, and B16F10 CAGA-dynGFP cells. Cells were challenged with TGM1-D12/TGM6 and TGFβ, and GFP signal after 21 hours was measured using IncuCyte (F). Schematic of the zHER2-link-TGM6 fusion protein. (**G**) Coomassie stain of SDS polyacrylamide gel to analyze the expression and purity of the produced zHER2-link-TGM6 fusion protein (Consecutive fractions 1 to 4 were pooled). (**H**) Effect of zHER2-link-TGM6 fusion protein on TGFβ-induced TGFβ/SMAD3-induced transcriptional reporter activity in wild-type NM18 or HER2 overexpression cells. zHER2-link-TGM6 (40 nM) was added 30 min prior to TGFβ (1 ng/ml, 15 hours) addition (**I**) Effect of zHER2-link-TGM6 fusion protein on TGFβ-induced SMAD2 phosphorylation TGFβ-induced stress fiber formation in wild-type or HER2 overexpression cells. zHER2-link-TGM6 (40 nM) was added 30 min before the addition of TGFβ (1 ng/ml). As a vehicle control elution buffer was used. (**J**) For SMAD2 phosphorylation and stress fiber formation, cells were treated for 1 hour or 2 days. Protein expression levels were analyzed by Western blotting. EMT was measured by F-actin staining.

Next, we examined the effects of hydroxychloroquine (an autophagy inhibitor that promotes lysosomal degradation), bafilomycin (a lysosomal inhibitor), and bortezomib and MG132 (proteasome inhibitors) on TGM6-induced reduction in TGFBR2 levels in NIH3T3 cells. We found that bafilomycin treatment blocked the TGM6-induced decrease in TGFBR2 expression (Figure 8D). In the presence of bafilomycin and absence of TGM6, we observed in NIH3T3 cells an increase in TGFBR2 levels (Figure 7D), consistent with the notion that TGFBR2 stability is controlled by lysosomal, but not proteasomal function [27]. Moreover, bafilomycin nearly completely rescued the inhibition of TGM6 on TGFβ-induced SMAD3 reporter activity (Figure 7E) and SMAD2 phosphorylation (Figure 7F) in NIH3T3 cells. Furthermore, co-treatment of NIH3T3 cells with TGM6 and cycloheximide, an inhibitor of protein synthesis, greatly potentiated the decrease in TGFBR2 levels compared to cycloheximide or TGM6 alone (Figure 7G). Taken together, our results reveal that in addition to competition between TGM6 and TGFβ for TGFBR2 (Figure 3B), TGM6 antagonizes TGFBR2 signaling by mediating LRP1-dependent TGFBR2 lysosomal degradation (Figure 7D-G).

### 2.10 Engineering TGM6-TGM1 chimeras to switch cell-type-specificity and change functionality

The modular structure of TGMs prompted us to generate TGM chimeras to determine whether we could alter cell selectivity and function. TGM1 mimics TGFβ’s effects on CD44-expressing cells, and TGM6 antagonizes TGFβ signaling in LRP1-expressing cells. We generated a chimera in which we switched TGM6-D4/5 for TGM1-D4/5 (termed TGM6-D3/TGM1-D4/5) (Figure 8A). Conditioned media of 293T cells transfected with an expression plasmid (or empty vector for control) were used to stimulate cells (Figure 8B). When TGM6 and TGM6-D3/TGM1-D4/5 were tested on NM18 or NM18 cells deficient in CD44, we found that TGM6-D3/TGM1-D4/5 mimicked TGM6 in inhibiting TGFβ signaling in control/wild-type NM18 cells but not in NM18 cells deficient in CD44 (Figure 8C). Whereas LRP1 is essential for TGM6 to inhibit TGFβ signaling in NM18 cells, TGM6-D3/TGM1-D4/5 inhibited equally TGFβ responses in wild-type/control or LRP1-deficient cells (Figure 8D). We thus changed the TGFβ antagonistic cell specificity of wild-type TGM6 for LRP^+^ cells with chimera TGM6-D3/TGM1-D4/5 to inhibit TGFβ signaling in CD44^+^ cells.

In another chimera, we fused TGM1 1D1/2 onto the N-terminus of TGM6 (termed TGM1-D1/2-TGM6) (Figure 8A). When this TGM chimera protein or TGFβ was used to challenge NM18 cells, we found that the chimera TGM1-D1/2-TGM6 and TGFβ potently activated the TGFβ/SMAD3 transcriptional response. We thus altered the TGM6 antagonist into a TGM1-D1/2-TGM6 agonist (Figure 8E).

Next, to explore the possibility of expanding the targeting of TGMs to fusions with an affibody/nanobody recognizing other cell surface receptors, we fused TGM6-D3 via a short linker sequence to an affibody [28] which engages with the human epidermal growth factor receptor 2 (HER2), and termed this zHER2-link-TGM6-D3 (Figure 8F). We expressed and purified the zHER2-link-TGM6-D3 protein (Figure 8G). Whereas zHER2-link-TGM6-D3 does not affect TGFβ-induced SMAD3 transcriptional response and SMAD2 phosphorylation in NM18 cells, these responses are blocked by zHER2-linkTGM6-D3 in NM18-HER2 overexpressing cells (Figure 8I). Furthermore, the TGFβ-induced morphological change of NM18 cells and stress fiber formation were not affected by zHER2-link-TGM6-D3 in wild-type cells but blocked in NM18-HER2 expressing cells (Figure 8J). Thus, the modular structure of TGMs enabled us to engineer synthetic TGM1/6 chimeras and fusions, thereby modulating cell-type specificity and functionality.

### 2.11. Engineering TGFBR2-VHH-based fusions for cell-type-specific targeting of TGFβ **signaling**

As the affinity of TGM6 D3 is much higher for mouse than human TGFBR2 ECD, the TGM6-D3-based fusion constructs are highly effective in inhibiting TGFβ signaling in mouse cells, but likely less so in human cells. Moreover, repeated *in vivo* administration of parasite-derived proteins into a mammalian host may elicit an immune response against the parasitic proteins. This may severely limit the potential of parasite-derived proteins as therapeutics due to their limited efficacy and safety. We therefore set out to design bispecific antibodies (BsAbs) without parasitic protein sequences that inhibit TGFβ signaling in a human cell-type-specific manner, similar to TGMs. We screened a phage display library for specific nanobody/variable heavy domain (VHH) binders to TGFBR2-overexpressed in HEK 293 T cells, and performed subsequent affinity maturation by yeast display using recombinant TGFBR2 ECD (Figure 9A). Specific TGFBR2-VHH binders were characterized by efficiency of immunoprecipitation of TGFBR2 and FACS analysis (Figure 9B, C). Clone 6 was identified as the strongest binder by immunoprecipitation (Figure 9B), and it was used in an affinity maturation assay to identify clone 6A as a high-affinity binder (Figure 9C). A binding assay revealed that the apparent 32.5 nM affinity for TGFBR2 to clone 6 was increased to 87 pM for clone 6A. Clone 6A had three mutations as compared to clone 6, one of these (Y62S) was solely responsible for the improved affinity and this single mutant was used in all further experiments. The effect of ectopic TGFBR2 overexpression (without exogenous added ligand on CAGA-GFP reporter activity) was slightly inhibited by clones 6A and 6Y625, but not by clone 6 (Figure S8A). Comparing the crystal structures of TGFBR2 with hTGFβ3 (PDB 1KTZ) and AlphaFold predicted hTGFBR2 ECD with TGFBR2-VHH revealed that the contact sites between the two protein pairs in the complexes overlap (Figure 9E). However, TGFBR2-VHH, when fused to Fc, was unable to inhibit TGFβ signaling (Figure 9F). For the purity assessment of TGFBR2-VHH-based proteins, see Fig S8B, C, and D. The biochemical and functional properties of TGFBR2-VHH are reminiscent of TGM6-D3, which, by itself, has high affinity for mTGFBR2 but cannot inhibit TGFβ signaling. We next fused TGFBR2-VHH to TGM1-D4/5 (Figure 9G) to see if this could regain TGFβ inhibitory activity in a cell type-specific manner. When tested for its effect on TGFβ-induced SMAD transcriptional response in wild-type or CD44-knockout NM18 cells, we found that TGFBR2-VHH-TGM1-D4/5 phenocopied the differential antagonistic effect of TGM6-D3-TGM1 D4/5 in wild-type versus CD44-depleted NM18 cells. Next, we fused TGFBR2-VHH to affibodies against HER2 (zHER2) (*28*) or EGFR (zEGFR) (*29*) (Fig 9G). These BsAbs were then tested on TGFβ-induced SMAD2 phosphorylation and SMAD3 transcriptional response in various TGFβ-responsive cell lines. Notably, EGFR2 and HER2 are selectively highly expressed in A431 lung adenocarcinoma and SKOV3 ovarian cancer cell lines, respectively (Figure 9H). Consistent with our expectation, zEGFR-TGFBR2-VHH and zHER2-TGFBR2-VHH inhibited TGFβ signaling selectively in A431 and SKOV3 cell lines, respectively (Figure 9J, K). Furthermore, similarly to what we found for zHER2-link-TGM6-D3, zHER2-TGFBR2 –VHH blocked TGFβ/SMAD-induced EMT in NM18-HER2 overexpressing, but not wild type NM18 cells (Figure 9L).

**Figure 9.**
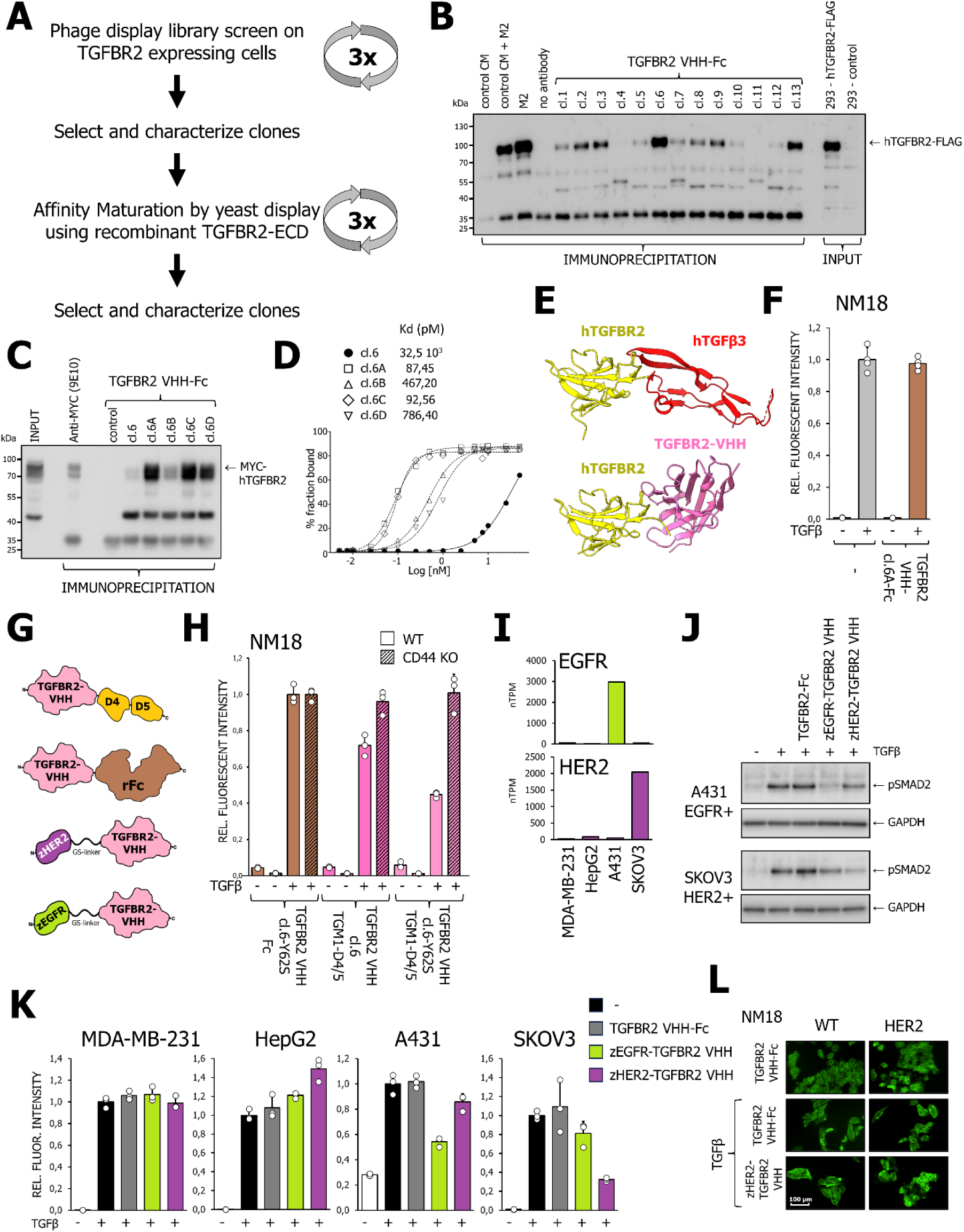
Designer TGFBR2 VHH-based bispecific antibodies that inhibit TGF-β receptor signaling in a cell-type-specific manner. (**A**) Schematic of the experimental flow that was followed for the identification of a high VHH nanobody recognizing TGFBR2 ECD with high affinity. (**B** and **C**) Analysis of the immunoprecipitation efficiency of TGFBR2 VHHs clones isolated by (**B**) phage display library screen using TGFBR2 overexpressing HEK 283 T cells and (**C**) affinity maturation by yeast display using recombinant TGFBR2 ECD. (**D**) Apparent affinity (Kd) measurements of various TGFBR2-VHH clones on TGFBR2 ECD. (**E**) Comparison of the crystal structure of hTGFβ3 to hTGFBR2 ECD (top panel) (PDB 1KTZ [66]) to the AlphaFold prediction of hTGFBR2 ECD in complex with TGFBR2-VHH-cl.6-Y62S. hTGFBR2 is in yellow, hTGFβ3 is in red, and TGFBR2-VHH is in magenta. (**F**) Effect of TGFBR-VHH cl.6 on TGFβ/SMAD3-induced transcriptional response (using CAGA-GFP transcriptional reporter) in NM18 cells stimulated with TGFβ (1 ng/ml). (**G**) Schematic presentations of TGFBR2-VHH-based protein fusions that were generated.; zHER2 and zEGFR are HER2 and EGFR high-affinity affibodies, respectively. (**H**) Effect of TGFBR VHH cl.6-Y62S fused to TGM1-D4/5 and control TGFBR2-VHH-cl.6-Y62Sfused to rFC on TGFβ/SMAD3 induced transcriptional response (using CAGA-GFP transcriptional reporter) in wild-type or CD44 knock-out NM18 cells stimulated with TGFβ (1 ng/ml). (**I**) Analysis of mRNA expression levels in nTPM (transcripts per million) of EGFR and HER2 in the indicated cancer cell lines using data from the Human Protein Atlas. (**J** and **K**) Effect of zEGFR-TGFBR2 VHH or zHER2-TGFBR2 VHH on TGFβ-induced (1 ng/ml) SMAD2 phosphorylation (**J**) and on TGFβ-induced SMAD3 transcriptional response (**K**) in the indicated cell lines. For vehicle controls, conditioned media were used. (**L**) Effect of zHER2-TGFBR2 VHH on TGFβ−induced EMT in NMuMG wild-type and HER2 overexpressing cells. The NM18 cells were stained with phalloidin-Alexa488 to visualize stress fibers.

## 3. Discussion

The murine helminth parasite-secreted TGM cytokines form a family of 10 structurally related proteins, of which TGM1, –4, and –6 have been shown to mimic or antagonize TGFβ-induced responses in selected host cell types [18–20, 29]. Here, we report the identification of TGM6 co-receptors. TGM6 directly partners with its co-receptor, LRP1, which is essential for TGM6’s antagonism and its *in-cis* cell-type-specific potency of TGFβ/SMAD signaling. In addition, TGM6, via its D3, interacts with co-receptor betaglycan, which attenuates the TGM6-mediated inhibitory effect on TGFβ/SMAD signaling. Thus, the two TGM6 co-receptors have an opposing function with respect to the modulation of TGFβ signaling. In line with what is expected for an antagonist, TGM6 binds with much higher affinity (15 nM) to mTGFBR2 than agonist TGM1 (0.6 μM) *(17)* and TGM4 (116 μM) [20]. TGM6 competes with TGFβ for TGFBR2 binding and induces TGFBR2 lysosomal degradation via LRP1. The modular structure of TGMs enabled us to generate TGM1/6 chimeras and inspired us to develop TGFBR2-VHH based BsAbs that modulate TGFβ signaling only in target co-receptor-expressing cells. We thus developed a toolbox of previously unavailable cell-type-restricted pharmacological manipulators of TGFβ/SMAD signaling, which can be expanded to other cell types based on the researcher’s needs.

A survey of mouse and human cell types from distinct tissues demonstrated that TGM6 could inhibit TGFβ signaling in mouse cells but not in human cells. As parasite TGMs have evolved through convergent evolution in mice, and not in humans, this provides a rationale for their mouse specificity. TGM6 interacts with TGFBR2 and LRP1 to antagonize TGFβ signaling. Our results indicate that TGM6 binding to LRP1 does not contribute to species-specific differences in TGM6 activity. We observed that (1) the ectopic expression of mLRP1 or hLRP1 was equally efficient in rescuing the TGM6 inhibitory effect in NIH3T3 TGFBR2 knock-out cells, and (2) that the mouse or human soluble LDLaIV cluster, i.e., the LRP1 interacting domain for TGM6, was equally effective in blocking the TGM6-induced inhibition of TGFβ signaling. Our results revealed that TGFBR2 is the critical determinant for TGM6 species specificity. Mouse, but not human, TGFBR2 rescued TGM6 antagonism in TGFBR2-knockout cells. ITC binding measurements revealed that TGM6-D3 bound mTGFBR2 with 40-fold greater affinity compared to hTGFBR2. Comparing the crystal structure of TGM6-D3:hTGFBR2 complex *(19)* with TGM6-D3:mTGFBR2 complex suggested that three residues, Phe47, Ser75, and Asp141, in mTGFBR2, corresponding to Leu47, Ala75, and Glu141 in hTGFBR2 ECDs, are the primary determinants underlying the species specificity for TGM6. This was confirmed by generating mTGFBR2 and hTGFBR mutants in which these residues were swapped, with residue 47 as the primary determinant.

Betaglycan was identified as a TGM6 co-receptor through a biased biochemical screen, in which TGM6 binding to the TGFβ family type 1, type II, and type III receptors was tested. Betaglycan is a cell-surface chondroitin and heparin sulfate protein of >300 kDa. It has an N-terminal orphan domain, a ZP domain, a transmembrane domain, and a short intracellular domain [31]. We found that for the TGM6-betaglycan interaction, both the orphan and ZP domains are required. This property is shared with TGFβ, for which the key TGFβ-binding regions in betaglycan in the orphan and ZP domain were recently identified [31]. While TGM6-D3 is sufficient and required for betaglycan interaction, betaglycan-TGM6 interaction is dependent on TGFBR2. Genetic depletion of betaglycan revealed that betaglycan is a negative regulator of TGM6’s antagonism of TGF-β/SMAD signaling. In the absence of betaglycan, TGM6 binding to LRP1, the mediator of TGM6’s antagonistic function, is slightly increased. The inhibitory function of betaglycan for TGM6 is in contrast to its role in TGF-β signaling, in which betaglycan aids in presentation of TGF-β to TGFBR and thereby potentiates TGFBR signaling [31, 32].

While TGM6-D3 has high affinity for mTGFBR2 (15 nM, Table 1) and TGM6 competes with TGFβ for TGBR2 binding on intact cells (Figure 3C), TGM6-D3 alone has no inhibitory effect on NIH3T3 cells [19]. Interestingly, we found that the absence of betaglycan results in TGM6 exhibiting inhibitory activity on TGF-β signaling at high TGM6 doses. Betaglycan may contribute to safeguard against weak TGM6 antagonism of TGFβ signaling in LRP1-deficient cells, which could be induced by TGM-D3-TGFBR2 interaction by TGM6.

Affinity labeling of cell surface proteins with iodinated TGM6 showed a prominent 600 kDa protein-TGM6 cross-linked complex. Mass spectrometric analysis of the pull-down of cell-surface proteins bound to TGM6 identified a 600 kDa protein, LRP1, as a putative interaction partner. These results were initially confirmed by the ability of the LRP1 antibody to specifically immunoprecipitate the TGM6-600 kDa protein complex. LRP1 is a multifunctional protein that plays an essential role in endocytosis and is recycled back to the membrane surface after internalization [26]. We mapped the TGM6-interacting domain on the LRP1 ECD to its LDLaIV cluster. TGM6-D4/5 binds to LRP1 and occurs independently of TGFBR2 and betaglycan. Genetic misexpression studies demonstrated that LRP1 is essential for the antagonistic function and cell-type-specific potency of TGM6. In both NIH3T3 and NM18 cells, LRP1 is required for TGM6 to antagonize TGFβ signaling. LRP1 is expressed at much lower levels in NM18 than in NIH3T3 cells. The negative regulator of TGM6, betaglycan, is, however, expressed at much lower levels in NM18 cells than in NIH3T3 cells and may explain why TGM6 remains a potent inhibitor of TGFβ signaling in NM18 cells. The modular structure of TGM6, with its D3 and D4/5 domains that independently engage TGFBR2/betaglycan and LRP1, respectively, represents an effective means of fine-tuning inhibitory effects across different cell types.

Notably, approximately 20 years ago, LRP1 was reported to act as a TGFβ receptor and was termed the TGFβ type V receptor [33–35]. It was shown to mediate TGFβ-induced inhibition of epithelial and myeloid cell proliferation. Other researchers have not followed up on this research.

TGFBR1 and TGFBR2 are both essential for TGFβ signaling. Our affinity-labeling immunoprecipitation studies demonstrated that TGM6 binds directly and with high affinity (15 nM) to mTGFBR2. TGM6 binds TGFBR2, mimicking the TGFβ-TGFBR2 interface [19]. TGM6 displaces TGFβ from interaction with TGFBR2 (and indirectly TGFBR1) on intact cells, in line with its antagonist function of TGFβ signaling. TGM6-LRP1 interaction may also indirectly contribute to TGM6 antagonism by promoting TGM6-TGFBR2 interaction through increased cellular avidity or by sequestering TGFBR1 from TGFBR2. Notably, these ligand-binding experiments were conducted with cells maintained on ice to prevent TGM6-induced receptor internalization. As mentioned earlier, high doses of TGM6-D3, in the absence of betaglycan, can inhibit TGF-β signaling. These results also suggest that competition between TGM6 and TGFβ for TGFBR2 contributes to TGM6’s antagonistic function. Importantly, we observed another mechanism by which TGM6 mediates its antagonism of TGFβ signaling. Challenging cells at 37°C with TGM6 potently decreased basal TGFBR2 expression, and the TGM6 co-receptor LRP1 was found to mediate TGFBR2 lysosomal degradation. Blocking TGM6-induced lysosomal degradation restored TGFBR2 levels and rescued TGFβ signaling in the presence of TGM6.

The modular structure of TGM6 may provide flexibility that allows it to interact not only with one cell type (and act *in-cis*) but with two adjacent cells (and act *in-trans*). However, experiments with mixed cultures of wild-type and LRP1-knockout cells did not support such *in-trans* signaling between neighboring cells. In addition, TGFBR2-expressing cells lacking LRP1 do not appear to function as a sink and do not inhibit the TGM6 effect on TGFBR^+^/LRP1^+^ cells. With LRP1 and betaglycan as co-receptors for TGM6, the expression levels of both will determine a cell’s responsiveness to TGM6-induced antagonism of TGFβ. Cells with high LRP1 and low betaglycan will be most responsive to TGM6.

Each TGM, characterized by its co-receptor specificity, serves as a powerful tool for investigating the function of TGF-β in specific cell types during mouse embryonic development, adult tissue homeostasis, and disease processes. Furthermore, as demonstrated by engineered TGM1/6 chimeras or fusions with the HER2 affibody sequence, we can create TGMs with varying cell-specificities and control their agonistic or antagonistic functions. D4/5 domains of TGMs can be exchanged with peptide-based engagers or nanobodies that specifically recognize certain co-receptors, thereby broadening the range of cell-specific TGFβ modulators. Furthermore, we developed a humanized TGFBR2-VHH that, by itself, has no effect on TGFβ signaling. However, when it is fused to TGM1 D4/5 or affibodies targeting EGFR or HER2, it enables cell-specific antagonism. While the cell-type specific mechanism of action is the same as TGM6-based antagonists, these TGFBR2-VHH-based BsAbs do not contain parasitic sequences and specifically target human cells.

## 4. Conclusion

The programmable pharmacological TGM-based modulators and TGFBR2-VHH-based BsAbs targeting TGFβ that are developed in our study are fundamentally different from current non-cell-selective TGFβ agonists and inhibitors [6]. The TGM-based engineering approach opens the possibility of creating a versatile set of designer agents to precisely manipulate TGFβ signaling tailored to specific mouse cell type(s). Moreover, these innovative TGFBR2-VHH-based BsAbs could serve as a foundation for developing new drugs targeting cancer and other human diseases associated with dysregulated TGFβ activity in a cell-selective manner, potentially avoiding the on-target side effects associated with existing TGFβ-targeting agents [6].

## 5. Experimental section

### 5.1. Materials

Cycloheximide (01810) and hydroxychloroquine (H0915) were obtained from Sigma-Aldrich; bafilomycin A1 (10-2060) was obtained from Focus Biomolecules; and bortezomib (SC-217785) was obtained from Santa Cruz. Recombinant human TGF-β3, expressed in *E. coli* as the mature C-terminal signaling domain, was refolded and purified by a patented method [36].

### 5.2. Cell lines

The following cell lines, 293T, NIH3T3, NM18 (a subline of the mouse mammary gland epithelial cell line NMuMG) [22], MDA-MB-231, B16F10, L6E9, HT-1080, MFB-F11, and HepG2 were maintained in DMEM (41966-052, Gibco), and U87-MG in EMEM (670086, Gibco), both supplemented with 10% fetal bovine serum (FBS) (S1810-500, Biowest) and penicillin/streptomycin. Generation of NIH-3T3 fibroblasts (CRL-1658) containing the fluorescent-based TGF-β/SMAD3 transcriptional reporter, i.e., CAGA-MLP-dynGFP, was previously described [37]. Cells were frequently tested for the absence of mycoplasma, and human cell lines were authenticated by Short Tandem Repeat (STR) profiling.

### 5.3. Selection, affinity maturation, and characterization of TGFBR2 VHH

TGFBR2 specific VHH’s were generated by Hybrigenics (Évry-Courcouronnes, France). A synthetic VHH library containing 3.109 clones was screened using HEK293T expressing TGFBR2. After three rounds of selection, 90 VHHs were picked and analyzed by Phage FACS for binding to TGFBR2; 28 were found to be positive. After sequencing and a confirmation analysis, we established 14 independent VHHs. The highest-affinity binder from our initial selection, i.e., clone (cl.) 6, was subsequently used for in vitro affinity maturation. To this end, a mutant TGFBR2-VHH library (2-5 × 10^6^) derived from the parental clone sequence was generated by introducing mutations restricted to the complementarity-determining regions (CDRs). Thereafter, the Yeast expressing mutant library was subjected to three rounds of sorting using recombinant carboxy-terminally avi-tagged TGFBR2 ECD as bait. The concentration of the TGFBR2 ECD domain was gradually decreased to select nanobodies with the highest binding affinities. Highest-affinity binders were characterized for the efficiency of TGFBR2 immunoprecipitation and FACS assays. All clones were DNA sequenced, and high-affinity clones, i.e., cl.6A to cl.6D, or mutational changes of different clones, i.e., cl.6-Y62S, were used in subsequent biological experiments. Sequences of the cl.6 and its derivatives are listed in Table S2. Other clone sequences are available upon request.

### 5.4. DNA constructs

#### LRP1 constructs

Full-length mLRP1 (non-tagged in pcDNA3) was a gift from Anton J. M. Roebroek, KU Leuven, Belgium [38]. A FLAG tag, along with an NheI site, was inserted between amino acids 23 and 24. PCR was used to generate deletion constructs in the pcDNA3 vector. The LDLaII and LDLaIV clusters were made by PCR and cloned into the pDISPLAY vector to create a membrane-bound protein. For stable expression, D4 and LDLaIV were cloned into a lentiviral vector (pLV-CMV-IRES-NEO). Cluster LDLaIV was also cloned from the pDISPLAY vector in pcDNA3 without the transmembrane domain (but with the signal peptide) to create a secreted protein. Full-length hLRP1 (without signal peptide and tag) in pCR8 (gateway donor vector) was a gift from Piet Gros, Utrecht University, the Netherlands [39]. Signal peptide was synthesized, and a full-length human LRP1 was constructed in pcDNA3. This construct was used to generate the D4 deletion mutant by PCR (in pLV-CMV-IRES-NEO). The full-length human LRP1 was used as a template to create a pDISPLAY construct with cluster LDLaIV (to make a membrane-bound protein. As with the mouse LDLaIV cluster, it was subcloned into pcDNA3, lacking the transmembrane domain (but retaining the signal peptide), to generate a secreted protein.

#### rBetaglycan (TGFβR3) constructs

rBetaglycan (TGFβR3) WT, delta Orphan, and delta ZP expression constructs were a gift from Fernando López-Casillas, Universidad Nacional Autónoma de México (UNAM), México City, México [40]. The constructs were re-cloned from the pCMV5 vector into a lentiviral vector (pLV-CMV-IRES-NEO).

#### TGFBR2 constructs

hTGFBR2 and mTGFBR2 were cloned into the pENTR1A vector. For mTGFBR2, a construct was generated in which four silent mutations were introduced in the target site of guide 2 to prevent it from being targeted. The GATEWAY system (Invitrogen) doxycycline-inducible lentiviral constructs were generated in the pCW57.1 vector, a gift from David Root (Addgene plasmid # 41393; http://n2t.net/addgene:41393; RRID:Addgene_41393

#### TGFBR2-VHH expression constructs

TGFBR2-VHH TGM1-D4-5, TGFBR2-VHH-Fc, zHER2-TGFBR2-VHH, and zEGFR-TGFBR2-VHH were made by cloning synthetic constructs into the pFUSE vector (Invivogen). For amino acid sequences, see Table S2.

#### HER2 expression construct

Human HER2 was cloned into the pLV-CMV-IRES-PURO by PCR from a HER2 cDNA clone (ccsbBroadEn_14631, Broad Institute).

#### Expression constructs for TGM1 and TGM6 (fusion) Proteins

Expression of *H. polygyrus* TGM1 and TGM6 (deletion) was conducted in HEK293 cells, using His-tagged proteins and metal-chelating affinity chromatography. Construction and purification methods are as previously described [8,9]. TGM6-D3/TGM1-D45 and TGM1-D12/TGM6 fusion constructs were generated by three-point ligation using two digested PCR products. The fragments were cloned into the SfiI and NotI sites of expression vector pSECTAG2a (Thermo Fisher Scientific). The TGM6-mut was generated by introducing mutations that replace Arg38, Ile78, and Tyr93 with alanine. Constructs were ordered from GeneArt, and *Asc*I and *Not*I restriction sites were used for insertion into pSecTag2A vector, and thereafter TGFM6-mut was expressed in HEK293 and purified as for other TGM6 proteins.

### 5.5. Expression and purification of TGM6-D3/TGM1-D45, TGM1-D12/TGM6 and TGFBR2-VHH based fusions

293T cells were transfected using PEI-MAX (Polysciences) with expression constructs for TGM6-D3/TGM1-D45, TGM1-D12/TGM6, and TGFBR2-VHH-based fusions using PEI-MAX® (24765, Polysciences). See Table S3 for information on the amino acid sequences of TGM6-D3/TGM1-D45m, TGM1-D12/TGM6, and TGFBR2-VHH-based fusion constructs. After 24 hours, the cells were washed 1x with phosphate-buffered saline (PBS), and fresh serum-free medium was added overnight. Next, the conditioned media were harvested, filtered through a 0.2 µm Puradisc FP 30 mm filter (Whatman), and used in the experiments. For vehicle control, conditioned medium from empty-vector-transfected cells was used. To compare TGM6-D3/TGM1-D45 with TGM6, TGM6 was also expressed in parallel and processed in the same manner.

### 5.6. Expression and purification of zHER2-link-TGM6-D3

zHER2-link-TGM6-D3 was produced using the ALiCE® system (AL00000003, LenioBio) using their pALiCE1 vector, all according to their manual. The proteins were purified using Strep-Tactin™ 4Flow™ beads (17577466, IBA Lifesciences) according to the manufacturer’s instructions. See Table S3 for information on the amino acid sequence of zHER2-link-TGM6-D3. The amino acid sequence and functional properties of zHER2-affibody were previously described [28].

### 5.7. Expression and purification of TGM6-D3 and TGFBR2 for ITC and crystallization

The amino acid sequences of the TGM6-D3, hTGFBR2, and mTGFBR2 proteins used in this study are presented in Table S4. TGM6-D3 and hTGFBR2 were each produced as insoluble inclusion bodies in E. coli BL21 (DE3), refolded, and purified by high-resolution ion-exchange chromatography (Source S and Source Q, respectively; Cytiva, Piscataway, NJ), as previously described [19, 41]. mTGFBR2 was produced and purified similarly to hTGFBR2, the only difference being that the final high-resolution ion-exchange purification using a Source Q column (Cytiva, Piscataway, NJ) was performed in 25 mM Tris, pH 8.0, instead of 25 mM MES, pH 6.0. The purity and identity of each protein were verified using non-reducing SDS-PAGE and by measuring the intact mass using liquid chromatography electrospray ionization time-of-flight mass spectroscopy (Micro TOF, Bruker Daltonics, Billerica, MA).

### 5.8. Isothermal titration calorimetry

ITC data were generated using a Microcal PEAQ-ITC instrument running version 1.40 of the Malvern PEAQ-ITC control software (Malvern Instruments, Westborough, MA). All experiments were performed in 25 mM HEPES, 150 mM NaCl, 0.05% NaN_3_, pH 7.4. Concentrations of the proteins in the syringe and sample cell are listed in Table 1. Before each experiment, all proteins were dialyzed into ITC buffer, then diluted as necessary before being loaded into the sample cell or syringe. For each titration, either 19 injections of 2.0 μL or 25 injections of 1.5 μL were performed, with an injection duration of 4 s and a spacing of 150 s; two independent data sets were collected for each titration. Integration and data fitting were performed using *Nitpic* 2.1.0 [42] and *Sedphat 15.2b* [23,24] with outliers removed as necessary. Each binding experiment, comprised of two independently measured titrations, was globally fit to a one-to-one binding model. The error estimates were derived using *SEDPHAT 15.2b* [23,24], and the data were plotted using *GUSSI 2.1.0* [43].

### 5.9. Crystallization, integration, reduction, and phasing

Crystallization was performed at 16 °C using the sitting-drop method in 96-well plates with 50 μL of well solution. The setup of crystallization screens was performed using a Mosquito robot (SPT Labtech, Hertfordshire, U.K.) by mixing 100 nL of protein complex with 100 nL of well solution. When looped, the crystals were transferred to a drop of well-solution adjusted with cryoprotectant. Each crystal was mounted in a nylon loop. Excess well solution was wicked off, and the looped crystals were flash-frozen in liquid nitrogen before being shipped at liquid nitrogen temperature for remote data collection. The TGM6-D3:mTGFBR2 complex crystal was crystallized with 35 mg mL^-1^ protein complex and a well-solution of 0.1M sodium HEPES pH 7.5, 10% PEG 4000, and 0.1M MgCl_2_. The cryoprotectant was 25% glycerol.

Diffraction data were collected at the Brookhaven National Lab National Synchrotron Light Source II (NSLS-II, beamline 17-ID-1). Diffraction images were integrated with iMosflm (*44*), and the space group of each crystal was confirmed with Pointless [45,46]. The diffraction data were reduced with Aimless [47], Ctruncate [48], and the Uniquify script [49] in the CCP4 software suite. Phaser [50] was used for molecular replacement, with the structure of the TGM6-D3:hTGFBR2 complex as the search model [19]. Several cycles of refinement using *phenix.refine* [51] and model building using COOT [52,53] were performed to determine the final structure. Data collection and refinement statistics are shown in the crystallography data table (Table S1). Images were generated using open-source PyMOL, with densities at 1.5 and 1.0, and root-mean-square deviation (RMSD) values of 0.5 and 0.4 for the human and mouse TGFBR2 complexes, respectively.

### 5.10. Guide RNA’s

Two different knockout guides against each target were designed using the ChopChop web tool (https://chopchop.cbu.uib.no/). The guides were cloned into the pLENTI-CRISPR-V2 plasmid (pLENTI-CRISPR-V2 was a gift from Feng Zhang, Addgene plasmid # 52961) [54]. See Table S5 for guide sequences.

### 5.11 Lentiviral cell transduction and selection

To generate stable cell lines, cells were transduced with a lentiviral vector produced in 293T cells. In brief, 293T cells were transfected (using PEI MAX® (24765, Polysciences)) with helper plasmids and the lentiviral plasmid, and on the following day, the transfection media were refreshed. On the second day, the conditioned media were collected and filtered to obtain a cell-free solution. Next, cells were incubated overnight with the filtered conditioned medium; 48 hours later, cells were selected with an appropriate antibiotic. After knockout selection, the knockout cells were used either as a pool or to generate single-cell clones.

### 5.12 Analysis of mRNA expression in cell lines

We analysed the mRNA expression in cancer cell lines using the Human Protein Atlas database (https://www.proteinatlas.org/) [55].

### 5.13 Transcriptional fluorescent protein-based reporter assay

To measure SMAD3 transcriptional activity, we used a fluorescence-based transcriptional reporter in which concatemerized SMAD3/4 binding elements (CAGA sequences) were cloned upstream of a minimal promoter. SMAD3/4 binding elements were derived from TGFβ/SMAD-responsive gene *SERPIN1*, encoding plasminogen activator protein 1 [36]. Cells containing either the CAGA-dynamic green fluorescent protein (dynGFP) or the CAGA-mCHERRYd2d2 reporter were seeded in 96-well plates. The next day, cells were pre-treated with TGM6 for 30 minutes and thereafter stimulated with TGFβ (in full media), and were placed in the IncuCyte S3 live-cell imaging analysis system (Sartoris). The cells were subsequently imaged at 3-hour intervals for 48 hours. Fluorescence intensity was analyzed using the IncuCyte software. TGF-β receptor/SMAD3 signaling was measured with an MFB-F11 cell assay as previously described [29]. All experiments were performed at least three times (and/or conducted on multiple independent cell clones), and representative results are shown.

### 5.14. Actin stress fiber staining

NM18 cells were grown in a 96-well dish at 20% confluence. Cells were pre-incubated with 100 ng/ml TGM6 for 30 minutes, after which the cells were stimulated with 1 ng/ml TGFβ for 48 hours. Cells were fixed with 4% paraformaldehyde (PFA) after 2 days, permeabilized with 0.1% Triton in PBS, blocked with 3% bovine serum albumin (BSA) in PBS, incubated with phalloidin-ALEXA-488 (Molecular Probes, A12379) for 1 hour, and then washed. The cells were imaged using the IncuCyte S3 (Sartorius) or a Leica. All experiments were performed at least three times, and representative results are shown.

### 5.15. Wound Healing assay

Cells were seeded in Image Lock plates (Sartorius, 4379) and were washed with PBS the next day. The cells were placed in DMEM containing 0,2% FBS and incubated overnight. Scratches were made using the Wound Maker (Sartorius). Cells were washed 1x with DMEM containing 0.2% serum. Cells were pre-incubated with or without TGM6 for 30 minutes and were subsequently stimulated with TGFβ or 10% FBS. Next, the cells were placed in the IncuCyte S3 live-cell imaging analysis system (Sartoris). The cells were imaged every hour for 1 day. Scratch sizes were analyzed using IncuCyte software (Sartorius). All experiments were performed at least three times, and representative results are shown.

### 5.16. Iodination and crosslinking

Purified recombinant proteins were labelled by iodination, biotinylation, and fluorescent coupling. Iodination of TGM1 proteins was performed by the chloramine T method, and cells were subsequently affinity-labeled with the radioactive ligand as previously described [56, 57]. In short, cells were incubated with the radioactive protein on ice for 3 hours. After incubation, the cells were washed, and crosslinking was performed using 0.27 µM disuccinimidyl suberate (DSS, Pierce, 21555) and 0.07 µM bis(sulfosuccinimidyl) suberate (BS3, Pierce, 21580) for 15 minutes. Cells were washed, scraped, and lysed. Lysates were incubated with antibodies overnight (4°C) and were precipitated using protein A Sepharose (Amersham, 17-0963-03). Samples were washed, boiled in SDS sample buffer, and subjected to SDS-PAGE. Gels were dried, exposed to a phosphor screen (FUJIFILM, BAS-SR2040), and then imaged using the Typhoon (Amersham). Antibodies used for immunoprecipitation are homemade in ten Dijke laboratory and have previously been described [57–60].

### 5.17. Western Blotting and detection

Cells were rinsed with ice-cold PBS before being lysed in lysis buffer. Protein concentrations were measured using the DC protein assay kit (Bio-Rad, 5000116), and samples were adjusted accordingly. After adding sample buffer and boiling for 5 minutes, the samples were separated on 8% SDS-polyacrylamide gels or on precast gradient gels (NuPAGE 4-12% gel, Invitrogen, WG1403BOX) and subsequently transferred to a nitrocellulose membrane (Protran, Amersham, 10600003). Transfer efficiency was assessed using Ponceau S. Membranes were blocked in TBS-T containing 5% skim milk for 1 hour at room temperature. Primary antibodies were incubated overnight at 4 °C. Secondary antibodies were incubated for 1 h at room temperature before detection using ECL (Clarity Western ECL Substrate, Bio-Rad, 170-5061) and the Chemidoc Imaging System (Bio-Rad). Band intensities were analyzed using Image Lab software (Bio-Rad). We used glyceraldehyde-3-phosphate dehydrogenase (GAPDH) expression or Ponceau S staining of the transferred protein as an equal loading control. See Table S6 for a list of the primary and secondary antibodies used in this study for Western blot analysis. All experiments were performed at least three times (and/or conducted on multiple independent cell clones), and results from representative experiments are shown.

### 5.18. Biotinylation and pulldown

Ten µg of TGM6 was incubated with EZ-Link™ Sulfo-NHS-LC-Biotin (Pierce, 21335) for 30 minutes at RT. The reaction was stopped by the addition of 50 mM Tris-HCl (pH 7.4). The reaction was purified using a G-25 column (GE Health, 28922529). For the pulldown, cells were washed with PBS and then incubated with biotinylated protein(s) for 3 hr on ice. After incubation, the plates were washed 3 times with PBS (Fresenius Kabi), and the cells were harvested in 1x Cell Lysis Buffer (Cell Signaling Technologies, 9803). After spinning, the supernatant was incubated with Neutravidin beads (Pierce, 29201) for 1 hr at 4°C (rotating). Beads were washed 4 times with lysis buffer and 3 times with 50 mM ammonium bicarbonate (Sigma-Aldrich, 09830), using fresh LoBind tubes for each wash (Eppendorf, 0030 108.116). The beads were resuspended in 250 µl of 50 mM ammonium bicarbonate with 250 ng of mass spec grade trypsin (Promega V5113), incubated overnight at 37°C (with agitation), after which peptides were recovered from the beads with a pre-washed 0.4 µm filter (Ultrafree MC HV, Millipore, UFC30HV00) and subjected to mass spectrometry analysis.

### 5.19. Mass spectrometry

Peptides were desalted using StageTips [61] and analyzed on an Orbitrap Fusion LUMOS (Thermo Fisher Scientific) hybrid mass spectrometer coupled to an EASY-nLC 1200 system (Proxeon, Odense, Denmark). Three technical repeats were performed, injecting 2%, 10%, and 50% of the sample, respectively. Digested peptides were separated using a 50 cm long fused silica emitter (FS360-75-15-N-5-C50, New Objective, Massachusetts, US) in-house packed with 1.9 μm C18-AQ beads (Reprospher-DE, Pur, Dr. Maisch, Ammerbuch-Entringen, Germany) and heated to 50°C in a Column Oven for ESI/Nano Spray (Sonation, Germany). Peptides were separated by liquid chromatography using a gradient from 2% to 32% acetonitrile with 0.1% formic acid for 30 minutes, followed by column reconditioning for 22 minutes. A Lock Mass of 445.12003 (polysiloxane) was used for internal calibration. Data were acquired in a Data-Dependent Acquisition (DDA) mode with a TopSpeed method, with a cycle time of 3 s, a scan range of 400-1500 m/z, and resolutions of 120,000 and 30,000 for MS1 and MS2, respectively, using the Orbitrap as detector in both cases. For MS2, an isolation window of 1.2 m/z and an HCD collision energy of 32% was applied. Precursors with a charge of 1 and higher than 6 were excluded from triggering MS2 as well as previously analyzed precursors with a dynamic exclusion window of 10s.

### 5.20. Mass spectrometry data analysis

Mass spectrometry data were analyzed using MaxQuant v1.6.14.0 [62] with the following modifications: the maximum number of missed cleavages by trypsin/p was set to 4. Searches were performed against an in silico-digested database of the mouse proteome, including isoforms and canonical proteins (Uniprot, 22nd April 2021). Oxidation (M), Acetyl (Protein N-canonical proteins (Uniprot, 22nd April 2021). Oxidation (M), Acetyl (Protein N-term), were set as variable modifications with a maximum of 3. Carbamidomethyl (C) was disabled as fixed modification. Label-free quantification was activated, not enabling Fast LFQ. The match-between-runs feature was activated with the default parameters.

### 5.21. Alphafold

The predicted structure of TGM6 with mTGFBR2 ECD and LRP1 mLRP1-LDLaIV was generated using AlphaFold3 (https://alphafoldserver.com/) [63].

### 5.22. Statistical analysis

Statistical analyses used one-way ANOVA or unpaired *t* tests, with correction for multiple comparisons, with Prism 9.

### 5.23. Scientific figure creation

Several figures from the NIAID Visual & Medical Arts (10/7/2024, NIAID NIH BioArt, (bioart.niaid.nih.gov)) and three figures from scidraw (scidraw.io, CC-BY 4.0) were used in the generation of the schematic figures 3A and 4A. Figures used from NIH bioart are: Receptor (BIOART-000429), Receptor protein (BIOART-000438), Generic Mass Spec Machine (BIOART-000639), Liquid Chromatography Machine (BIOART-000638), and Mass Spectrometry Graph (BIOART-000582). Figures used from scidraw; Eppendorf tube closed and Petri Dish by Diogo Losch De Oliveira, and Progenitors by Roberta Schellino.

## Acknowledgments

We thank Midory Thorikay for expert technical assistance.

## Author contributions

PtD, AH, RM, and MvD conceived and designed the study. MvD, TS, KF, JZ, GvdZ, LP, CH, CC, AM, RG-P, and PvV designed and performed the experiments. MvD, TS, JZ, KF, GvdZ, LP, CH, AM, R G-P, PvV, RMM, APH, and PtD analyzed the data. PtD coordinated the study and wrote the paper with input from all authors. RMM, PtD, JZ, KF, and APH provided funding.

## Ethics statement

No clinical samples were used, nor were animal studies performed.

## Data and materials availability

All data needed to evaluate the conclusions in the paper are present in the paper and/or the Supplementary Materials. All unique/stable reagents generated in this study are available from the lead contact upon reasonable request.

## Permission to reproduce materials from other sources

We have not used material from other sources.

## Declaration of generative AI and AI-assisted technologies in the writing process

During the preparation of this work, the author(s) used Grammarly to check grammar and spelling. After using this tool/service, the author(s) reviewed and edited the content as needed and take(s) full responsibility for the content of the published article.

## Resource availability

### Lead contact

Requests for further information and resources should be directed to and will be fulfilled by the lead contact, Peter ten Dijke.

## Funding

This work was funded by grants awarded from the Oncode Institute base fund and NWO (Grant ID: https://doi.org/10.61686/AHJBX34229) to PtD, Chinese Scholarship Council (to JZ and KF), a Wellcome Trust Discovery award to RMM, APH, and PtD, Wellcome Trust grant (306173) to RM, and NIH R03 (AI53915) and F30 (AI157069) grants to APH and AM, respectively.

## Notes

### Competing Interest Statement

Peter ten Dijke, Andy Hinck and Maarten van Dinther have applied for a patent on cell-type specific TGFbeta inhibitors. All other co-authors have no conflict of interest.

