## Supplemental information for "Decoding the Mechanism of Action of a Parasite TGFβ antagonist Inspires the Creation of Cell-type-specific TGFβ Modulators"

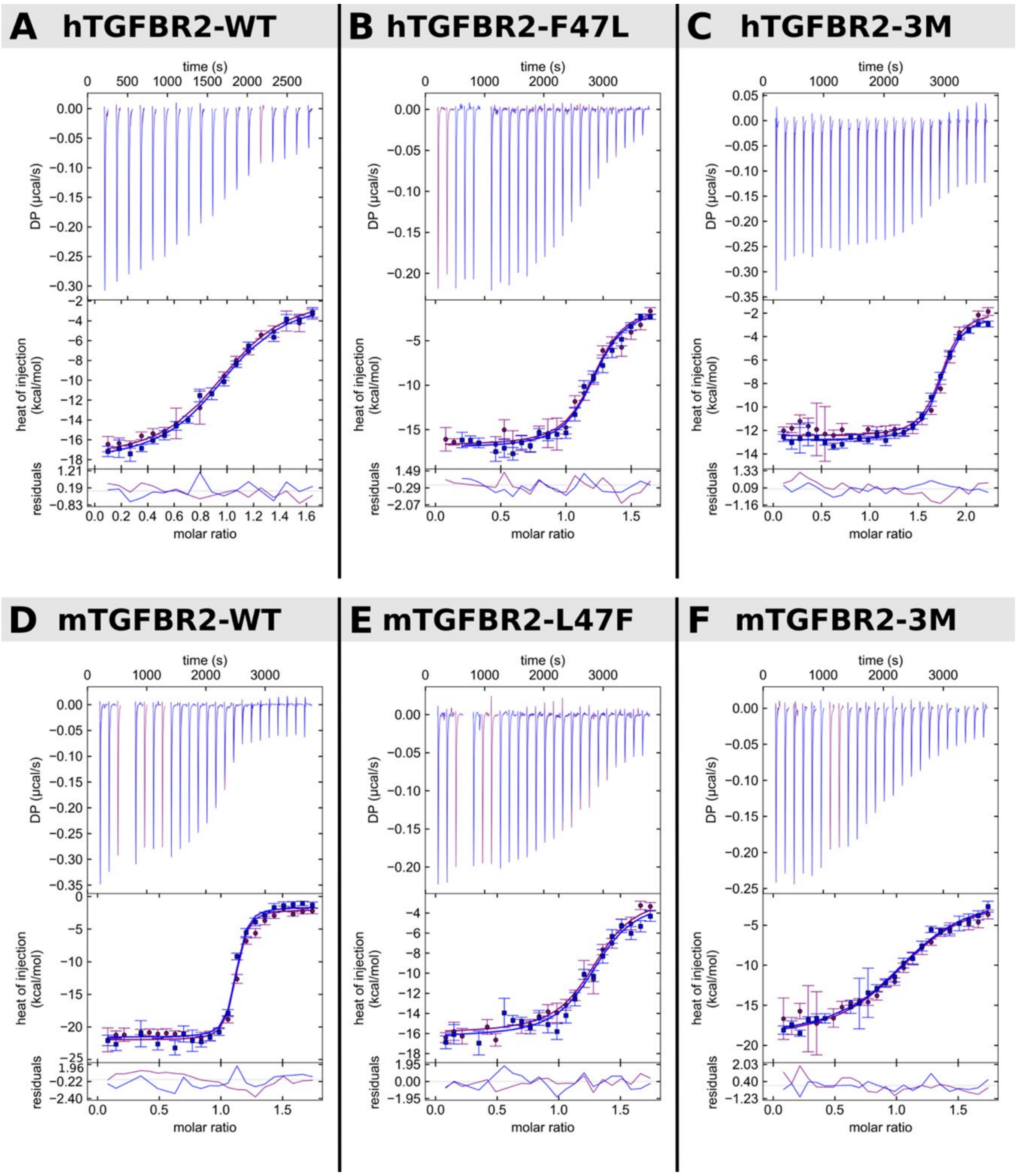


**Figure S1. Isothermal titration calorimetry data for binding of TGM6-D3 to wild-type mouse and human TGFBR2 ECD and variants**. (A-F) ITC binding data between TGM6-D3 in the cell and TGFBR2 in the syringe. Wild-type human TGFBR2 and the F47L and F47L, S75A, D141E (3M) variants are shown in panels A-C, respectively, while wild-type mouse TGFBR2 and the L47F and L47F, A75S, and E141D (3M) variants are shown in panels D-F, respectively. Each ITC sub-panel comprises thermograms (upper panel), mean integrated heats (middle panel), and fit residuals (lower panel). Duplicate datasets were collected (purple and blue), and the data from both were globally fit to a 1:1 model to obtain a single KD, DH, and incompetent fraction in the cell and syringe (shown by continuous lines in the middle panel). Fitted values are provided in Table 1.

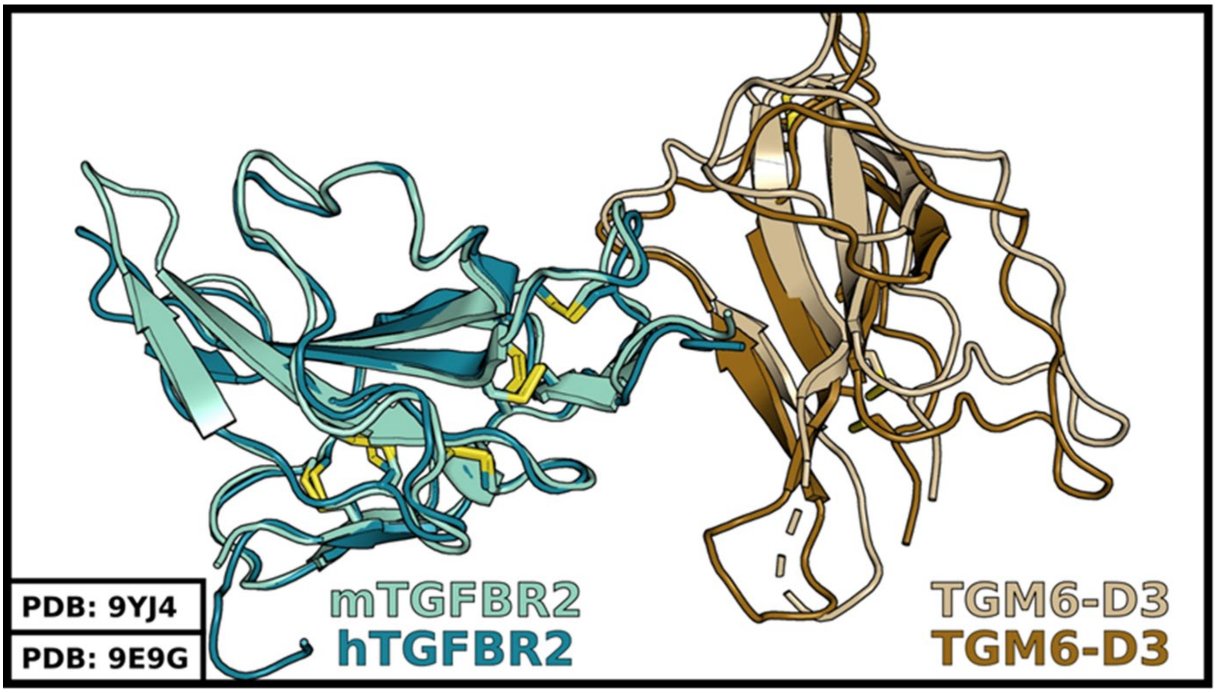


**Figure S2. Overall Structure of TGM6-D3 in complex with hTGFBR2 or mTGFBR2.** Asymmetric unit of TGM6-D3 in complex with hTGFBR2 has a single copy of the complex. Overlay of mTGFBR2 (light green):TGM6-D3 (light brown) and hTGFBR2 (dark green):TGM6-D3 (dark brown).

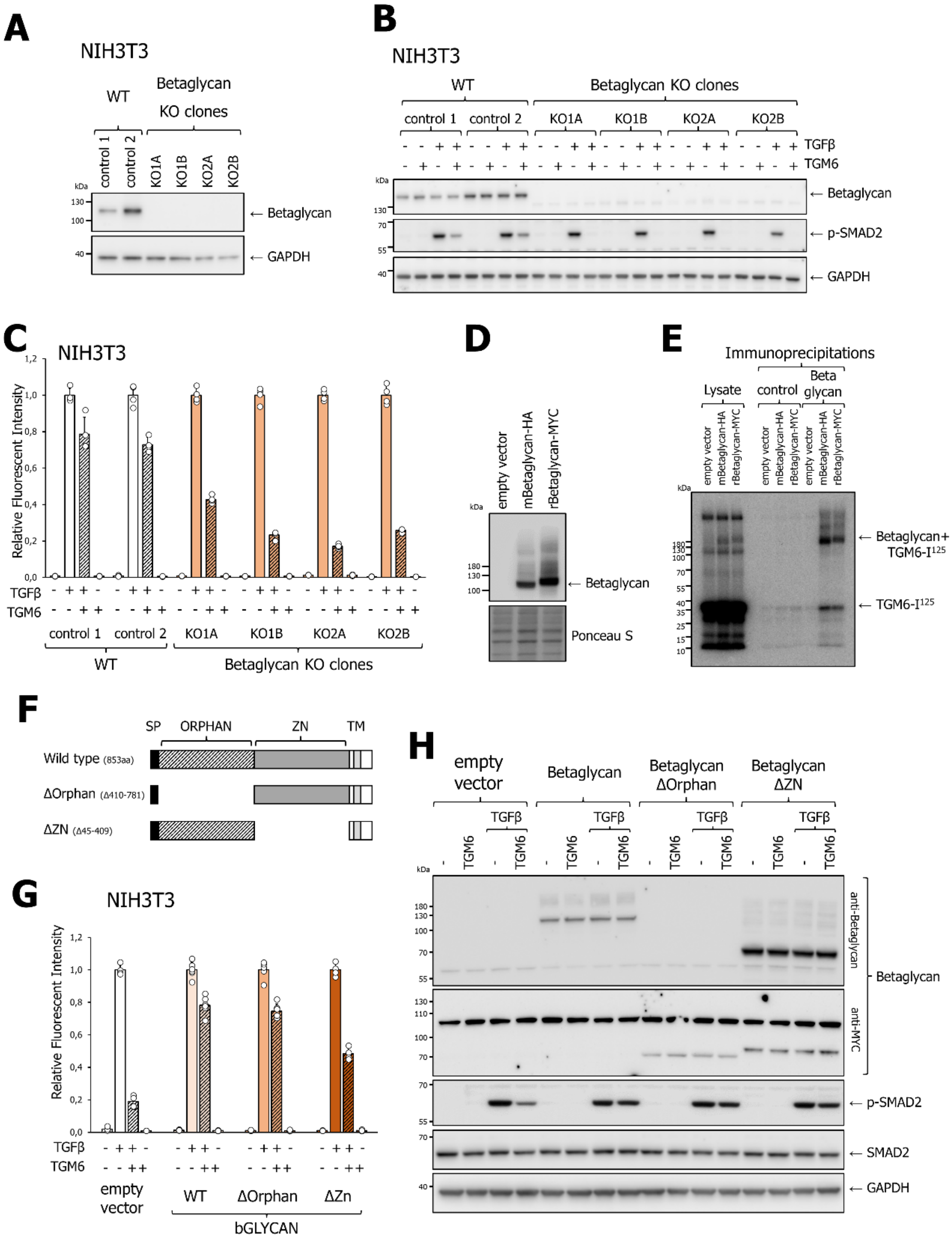


**Figure S3 Betaglycan is negative regulator of TGM6 antagonism of TGFβ/SMAD signaling response and Orphan and Zona Pellucida domain of Betaglycan are both functionally required.**

(A) Analysis of betaglycan expression in wild-type (control 1 and 2) and knock out clones (KO1, KO2, KO3 and KO4) as measured by Western blot analysis. (B and C) Effect of betaglycan knockout on antagonism by TGM6 on TGFβ-induced SMAD2 phosphorylation (B) and TGFβ/SMAD3 transcriptional reporter response (C).(D) Analysis of ectopic expression of mouse and rat betaglycan in L6E9 myoblasts. Mouse or rat betaglycan was transiently expressed in L6E9 cells. (E) Betaglycan-transfected cells (characterized in A) were used to interrogate the  binding of TGM6 to mouse or rat betaglycan that were transiently expressed in L6E9 cells. Iodinated TGM6 was used to affinity-label cell-surface proteins; betaglycan was immunoprecipitated from cell lysates, and signals were analyzed by autoradiography. (F) Schematic representation of two betaglycan deletion constructs. (G and H) Effect of wild-type betaglycan and betaglycan deletion mutants in L6E9 myoblasts on TGFβ/SMAD3-induced dynGFP transcriptional response (G) and phosphorylation of SMAD2 (H). TGM6 (100 ng/ml) was added 30 min before the addition of TGFβ (1 ng/ml). TGFβ treatment induced a transcriptional response and phosphorylation of SMAD2 at 21 hours and 1 hour, respectively. CAGA-dynGFP was measured by IncuCyte live-cell imaging (G), and protein levels were assessed by Western blot analysis (H). Fig. S3B and S3C. correspond to the extended dataset for Figs. 3D and 3E.

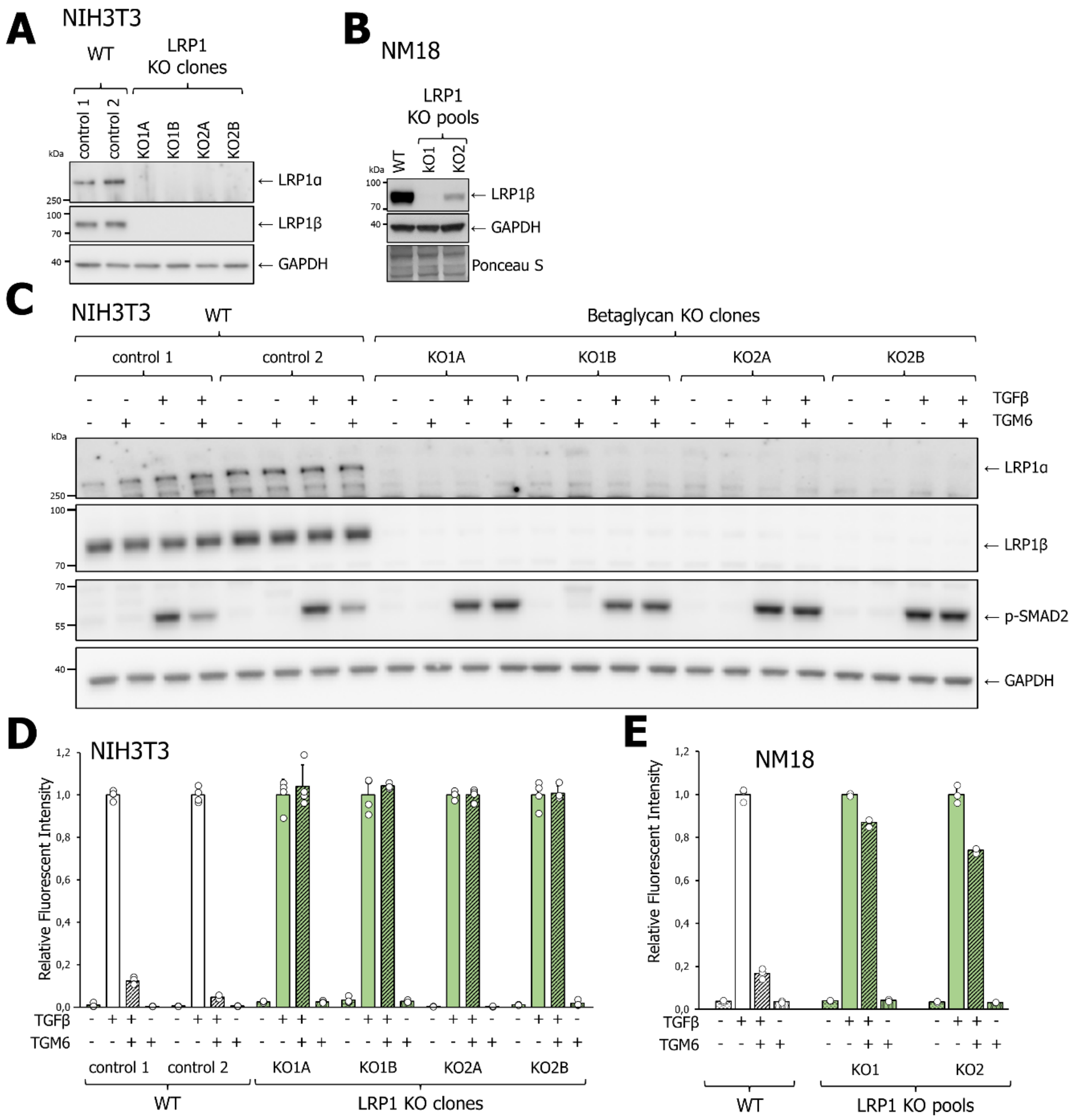


**Figure S4 LRP1 is a co-receptor for TGM6**

(A) and (B) Validation of LRP1 knock-out in NIH3T3 and NM18 cells. Expression of LRP1 (sub-units LRP1α and LRP1β) as measured by Western blot analysis in NIH3T3 wild-type clones (control 1 and control 2) and LRP1 KO clones (KO1A, KO1B, KO2A, and KO2B). KO1 and KO2 were made using different guide RNAs. (B) Expression analysis of LRP1β in LRP1 wild-type cells and LRP1 KO pools (KO1 and KO2) as measured by Western blot analysis. (C) Effect of LRP1 deficiency in NIH3T3 cells on antagonism of TGM6 on TGFβ/SMAD2 phosphorylation response (C) and (D and E) on the antagonism of TGM6 on TGFβ/SMAD3 transcriptional activity in NIH 3T3 and NM18 cells. For NIH3T3, the results of two controls (control 1 and 2) and 4 KO clones (KO1A, KO1B, KO2A, KO2B) are shown, and for NM18 th results of wt and two LRP1 knockout pools (KO1 and KO2) are shown. Fig. S4C, S4D, S4E correspond to the extended data set for Fig. 4C, 4D, and 4E.

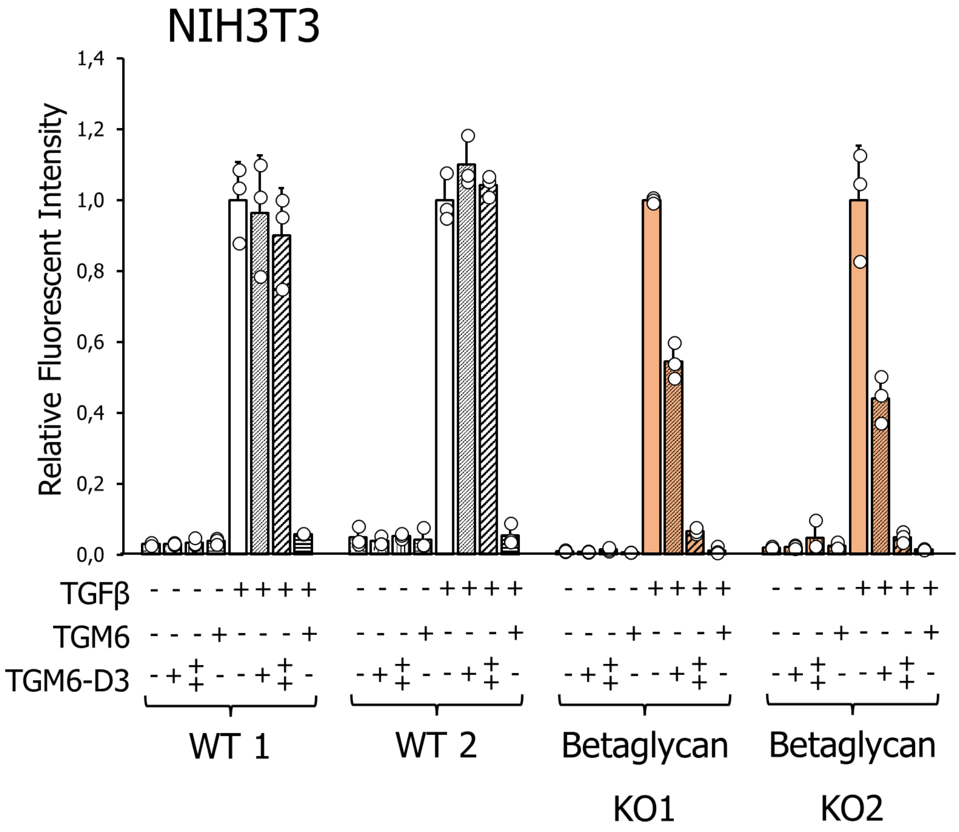


**Figure S5. Effect of betaglycan deficiency on the ability of TGM6-D3 to antagonize TGFβ signaling.**

Effect of betaglycan deficiency on the ability of TGM6-D3 to antagonize TGFβ-induced CAGA-dynGFP reporter activity in NIH3T3 control clones, and betaglycan NIH3T3 KO clones. Cells were pre-incubated with TGM6 (100 ng/ml) or TGM6-D3 (1000 or 5000 ng/ml) for 30 min before stimulation with 1 ng/ml TGFβ for 21 h. The results on two wild-type clones (Wt1 and WT2) and two Betaglycan KO clones (KO1 and KO2) are shown. KO1 and KO2 were made using different guide RNAs. Fig. S5 corresponds to the extended dataset for Fig. 5H.

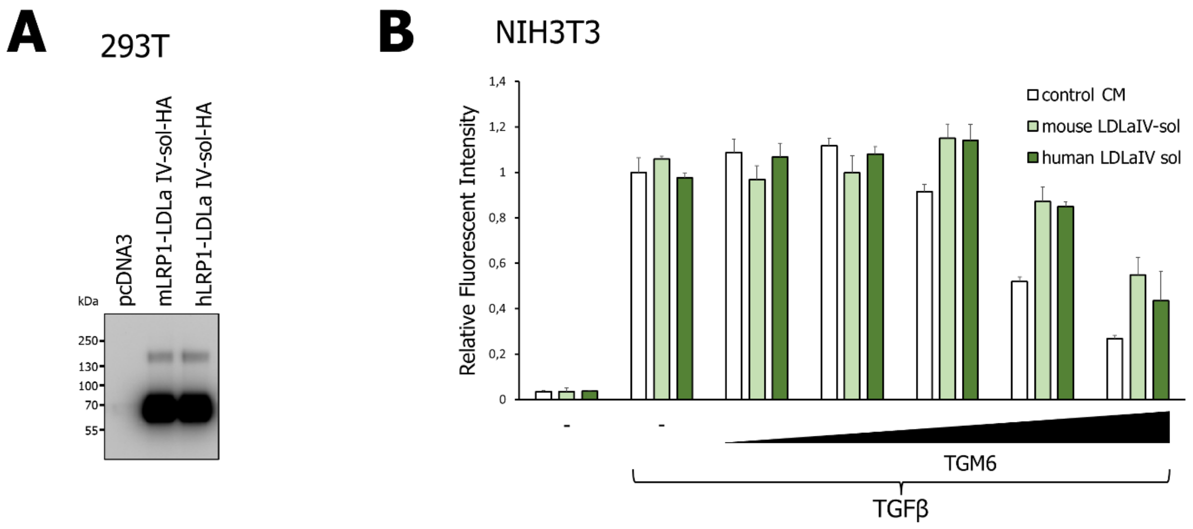


**Figure 6S Effect of soluble LRP1-LDLaIV on TGFβ/SMAD signaling**

(A) Expression analysis of soluble mLRP1-LDLaIV or hLRP1-LDLaIV by western blot analysis of conditioned media of transfected HEK293T cells. (B) Effect of mLRP1-LDLaIV or hLRP1-LDLaIV on TGM6-mediated antagonism of TGFβ/SMAD3-induced transcriptional response in NM18 cells. TGM6 (2,5, 10, 25, 50, or 100 ng/ml) was pre-incubated with conditioned media containing either mLRP1-LDLaIV or hLRP1-LDLaIV. Subsequently, this was used to pre-incubate NM18 cells (30 min) before stimulation with 1 ng/ml TGFβ for 21 h.

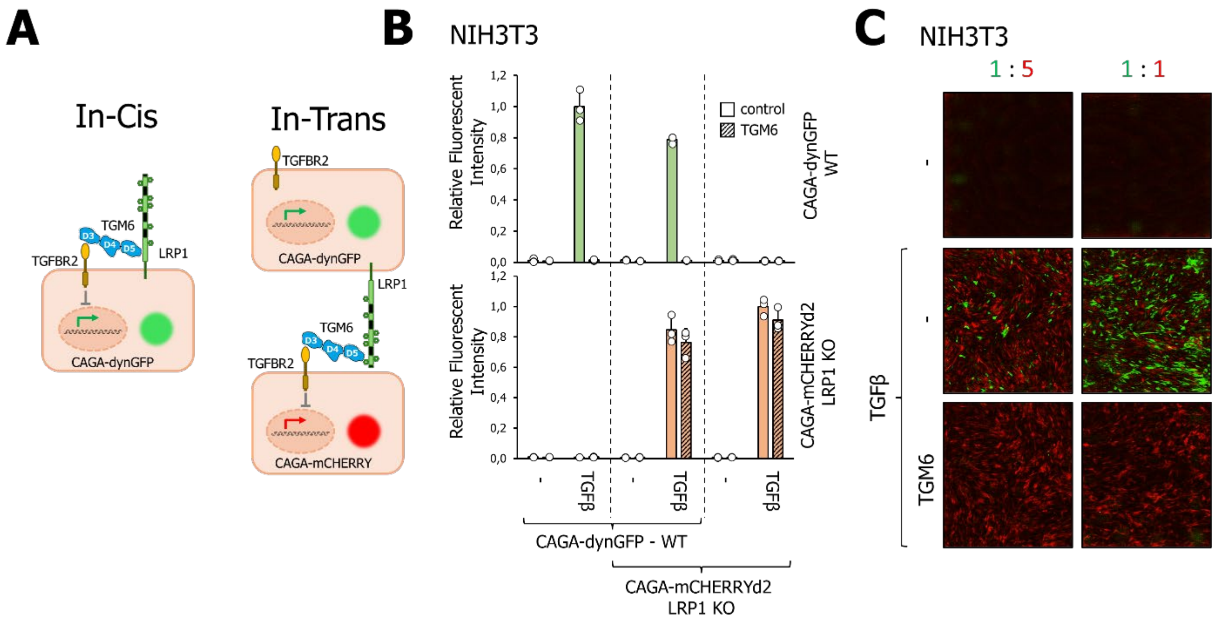


**Figure S7 TGM6 elicits cellular effects in-cis and not in-trans**.
(A) Schematic representation of TGM6 signaling in-cis or in-trans. (B and C) Effect of TGM6 on NIH3T3-CAGA-dynGFP cells (TGFBR2+LRP1+) and NIH3T3-CAGA-mCHERRYd2 cells deficient in LRP1 (TGFBR2+LRP1-), either as mono- or mixed cultures in two different ratios. Cells were challenged with 100 ng/ml TGM6 and/or 1 ng/ml TGFβ, and CAGA transcriptional response was measured after 21 hours. Quantification is shown in B, and some representative fluorescent images are shown in C.

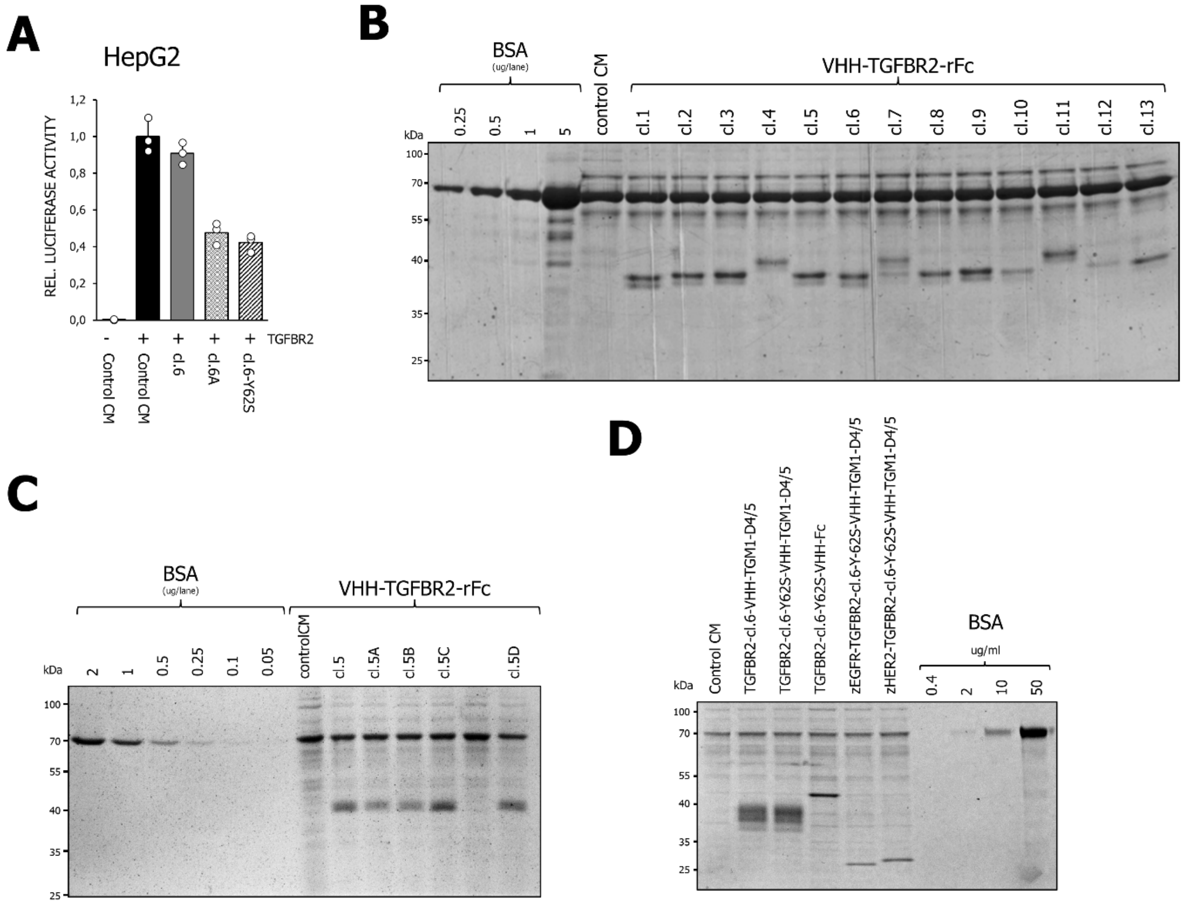


**Figure S8 Expression controls fusion protein and reporter assay**
(A) CAGA-luciferase transcriptional reporter assay in HepG2 cells showing the effect of various TGFBR2 VHH's on TGFBR2 overexpression induced signaling. (B, C and D) Expression controls of the various TGFBR2-based fusion proteins used in Fig. 9. Different amounts of bovine serum albumin (BSA) is used to estimate protein concentrations.
